# A vast world of viroid-like circular RNAs revealed by mining metatranscriptomes

**DOI:** 10.1101/2022.07.19.500677

**Authors:** Benjamin D. Lee, Uri Neri, Simon Roux, Yuri I. Wolf, Antonio Pedro Camargo, Mart Krupovic, RNA Virus Discovery Consortium, Peter Simmonds, Nikos Kyrpides, Uri Gophna, Valerian V. Dolja, Eugene V. Koonin

**Author notes:** Correspondence: Eugene V. Koonin < >.

## Abstract

Viroids and viroid-like agents are unique, minimal RNA replicators that typically encode no proteins and hijack cellular enzymes for their genome replication. As the extent and diversity of viroid-like agents are poorly understood, we developed a computational pipeline to identify viroid-like covalently closed circular (ccc) RNAs and applied it to 5,131 global metatranscriptomes and 1,344 plant transcriptomes. The search resulted in 11,420 viroid-like, ribozyme-containing cccRNAs spanning 4,409 species-level clusters, which is a five-fold increase compared to the previously known set of viroids and viroid-like RNA agents. Within this diverse collection, we identified numerous putative novel viroids, satellite RNAs, retrozymes, and ribozylike viruses. We also found previously unknown ribozyme combinations and unusual ribozymes within the cccRNAs. Self-cleaving ribozymes were identified in both RNA strands of ambiviruses and some mito-like viruses as well as in capsid-encoding satellite virus-like cccRNAs. The broad presence of viroid-like cccRNAs in diverse transcriptomes and ecosystems implies that their host range is not limited to plants, and matches between viroid-like cccRNAs and CRISPR spacers suggest that some of them might replicate in prokaryotes.

## Introduction

Viroids, which cause several economically important diseases in agricultural plants, are the smallest and simplest among the known infectious agents (Daròs et al., 2006; Diener, 2001; Mascia and Gallitelli, 2017). Viroids are small, covalently closed circular (ccc) RNA molecules of 220 to 450 nucleotides that encode no proteins and consist largely of RNA structures that are required for replication or viroid-host interaction. In contrast to viruses, which hijack the host translation system to produce proteins encoded in virus genes, viroids take advantage of the host transcriptional machinery. Specifically, viroids hijack the host plant’s DNA-dependent RNA polymerase II to transcribe their RNA and thus catalyze viroid replication (Mühlbach and Sänger, 1979; Navarro et al., 2000; Schindler and Mühlbach, 1992). Viroids utilize the rolling circle replication (RCR) mechanism, producing multimeric intermediates that are cleaved into genome-size monomers by ribozymes that are present in both the plus and the minus strands of viroid RNA or by recruited host RNAses (Branch and Robertson, 1984; Flores et al., 2017). The resulting linear monomers are then ligated by a host DNA ligase to form the mature cccRNA (Nohales et al., 2012a, 2012b).

Since the discovery of viroids in 1971 (Diener, 1971), about fifty distinct viroid species have been identified in plants and classified into two families, *Avsunviroidae* and *Pospiviroidae*. Members of the *Avsunviroidae* use viroid-encoded autocatalytic hammerhead (HHR) ribozymes, a defining feature of this family, to process replication intermediates into unit length viroid genomes (Di Serio et al., 2017; Wang, 2021). Members of the family *Pospiviroidae* lack ribozymes and instead rely on conserved sequence motifs that serve as recognition and cleavage sites for host RNase III (Branch et al., 1988). The members of the two viroid families also adopt distinct RNA structures: a branched RNA conformation is predominant in avsunviroids (Giguère et al., 2014a), in contrast to the typically rod-shaped conformation of pospiviroids (Giguère et al., 2014b).

In addition to viroids, several other groups of infectious agents also possess genomes consisting of cccRNA (de la Peña et al., 2020). Many plant viruses support the replication of small (about 300 nt) circular satellite RNAs (sometimes called virusoids but hereafter, satRNAs) that closely resemble viroids (Navarro et al., 2017) and also replicate via the rolling circle mechanism (Bruening et al., 1991). The satRNAs differ from viroids in that they are replicated by the RNA-dependent RNA polymerase (RdRP) of the helper virus and are encapsidated in that virus’s capsid (Huang et al., 2017; Rao and Kalantidis, 2015). Thus, these satRNAs are effectively encapsidated viroids. Unlike viroids, satRNAs encode both HHRs and hairpin ribozymes (HPR), a distinct ribozyme variety (Ferré-D’Amaré and Scott, 2010).

Another notable viroid-like agent is the so-called retroviroid, carnation small viroid-like RNA (CarSV) which, unlike viroids, does not appear to transmit horizontally among plants (Daròs and Flores, 1995). CarSV, the only currently known retroviroid, is a cccRNA that is similar to viroids in size and contains HHRs in both strands. However, in contrast to the viroids, an extrachromosomal DNA copy of CarSV has been discovered and shown to integrate into the plant genome with the help of a pararetrovirus (Hegedus et al., 2004; Vera et al., 2000).

A recently discovered distinct group of cccRNA agents are retrozymes, retrotransposons that propagate via circular RNA intermediates of about 170 to 400 nucleotides. The retrozymes are viroid-like in that they do not encode any proteins but contain self-cleaving HHRs (Cervera et al., 2016; de la Peña and Cervera, 2017). However, unlike viroids, the retrozymes are neither infectious nor autonomous, but rather, hijack the replication machinery of autonomous retrotransposons. Resembling satRNAs and avsunviroids, retrozyme cccRNAs also adopt a branched conformation.

A distinct group of viroid-like agents is the viral realm *Ribozyviria* (Hepojoki et al., 2020) that includes deltaviruses, such as hepatitis delta virus (HDV), an important human pathogen. Similarly to pospiviroids, ribozyviruses possess rod-shaped cccRNA genomes that replicate via the rolling circle mechanism and encode distinct ribozymes, unrelated to those of viroids, that autocatalytically process multimeric replication intermediates (Kos et al., 1986; Modahl et al., 2000; Sureau and Negro, 2016). Ribozyviruses have substantially larger genomes than viroids (about 1.7 kb), encode their own nucleocapsid protein, and rely for reproduction on a helper virus (hepatitis B virus in the case of HDV), which provides the envelope protein for ribozyvirus virions. For years, HDV remained the only known deltavirus. Recently, however, viruses more distantly related to HDV have been discovered in diverse vertebrates and invertebrates including rodents (Paraskevopoulou et al., 2020), bats (Bergner et al., 2021), snakes (Hetzel et al., 2019), birds (Wille et al., 2018), fish (Chang et al., 2019) and termites (Chang et al., 2019), suggesting a considerable uncharacterized diversity of ribozyviruses.

Viroids and viroid-like cccRNAs comprise a fundamentally distinct type of minimal replicators, or ultimate parasites, that lack genes and effectively consist only of RNA structures required for replication. This extreme simplicity of viroids triggered speculation on a potential direct descendance of viroids from primordial RNA replicators (Diener, 2016; Flores et al., 2022). However, the apparent narrow host range of viroids, which so far have been reported only in plants, does not appear to be easily compatible with such an evolutionary scenario. Instead, given the major similarities between retrozymes and avsunviroids, it has been suggested that avsunviroids descended from retrozymes (de la Peña and Cervera, 2017; Lee and Koonin, 2022).

Given the ultimate structural simplicity of viroids and related cccRNAs and the universality of the DNA-dependent RNA polymerases involved in their replication across life forms, the current narrow spread and limited diversity of parasitic cccRNAs appear puzzling. Furthermore, this apparent paucity of viroid-like agents is in stark contrast to the burgeoning diversity of RNA viruses, many thousands of which including numerous, distinct new groups have been discovered by metatranscriptome analyses (Edgar et al., 2022; Neri et al., 2022; Wolf et al., 2020; Zayed et al., 2022). At present, there are at least three orders of magnitude more known RNA viruses than there are viroids and viroid-like cccRNAs.

We were interested in investigating the global diversity of viroids and viroid-like agents. To this end, we performed an exhaustive search for cccRNAs in a collection of about 5,131 diverse metatranscriptomes that have been recently employed for massive RNA virus discovery (Neri et al., 2022), and additionally searched 1,341 plant transcriptomes (One Thousand Plant Transcriptomes Initiative, 2019). This search yielded 10,183,455 putative cccRNAs within the size range of 100–10,000 nt, of which 11,378 were classified as viroid-like based on the presence of predicted self-cleaving ribozymes or direct similarity to reference sequences. This set of cccRNAs represents an about five-fold increase of the known diversity of viroid-like agents. Further analysis of these cccRNAs led to the identification of numerous putative novel viroids, satRNAs, retrozymes and ribozy-like viruses.

## Results

### Computational approach for the discovery of viroid-like cccRNAs

We first developed an integrated, scalable computational pipeline for the *de novo* discovery and analysis of viroids and viroid-like cccRNAs directly from assembled transcriptomes and metatranscriptomes (Figure 1). The pipeline starts with the reference-free and *de novo* identification of cccRNAs or RCR intermediates. This method depends on the fact both complete circular monomers and multimeric linear intermediates will be assembled containing detectable head-to-tail repeats (Qin et al., 2020). Once identfied, the sequences are then cleaved to unit length and deduplicated, taking circularity into account. Starting from the set of detected cccRNAs, the pipeline performs both alignment-free and alignment-based searches. The primary approach for the identification of viroid-like agents among the cc-cRNAs is the prediction of self-cleaving ribozymes using RNA sequence and secondary structure covariance models (Nawrocki and Eddy, 2013). Under the premise that the diversity of ribozymes in viroid-like RNAs could be greater than so far uncovered, we curated a database of known self-cleaving ribozyme models from Rfam (Kalvari et al., 2021), of which only a minority have been described in viroids and viroid-like RNAs. We supplemented this model database with the pospiviroid RY motif (Gozmanova, 2003) to enable the search to detect potential pospiviroids, which do not contain ribozymes.

**Figure 1:**
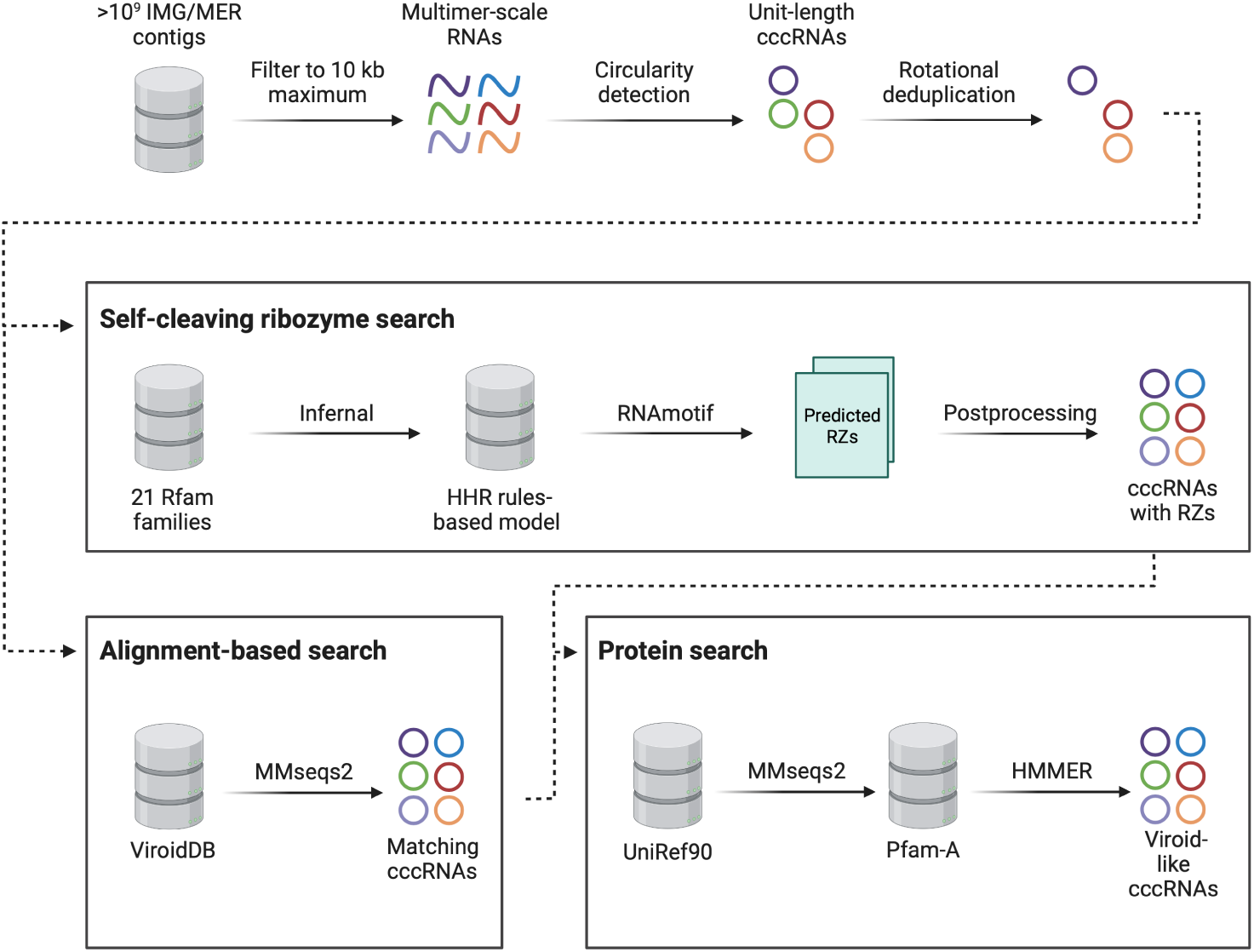
Viroid-like cccRNA detection pipeline.

The pipeline also performs direct sequence similarity searches against reference databases such as ViroidDB (Lee et al., 2021).

Ribozyme-containing cccRNA sequences were classified as symmetric or asymmetric depending on whether they contained predicted ribozymes in both or only one RNA polarity, respectively, reflecting the RCR mode these cccRNAs are likely to undergo. However, it cannot be ruled out that some apparently asymmetric cccRNAs actually contain a second ribozyme distinct from the currently known ones.

We validated this method by demonstrating its ability to recover known viroid-like RNAs in both transcriptomes and metatranscriptomes (Table S1). For the transcriptomic validation, we processed and searched the 1,000 Plant transcriptome (1KP) data set (One Thousand Plant Transcriptomes Initiative, 2019). We chose this data set due to the known presence of all type of viroid-like cccRNAs except ribozyviruses in plant transcriptomes. Assembling the raw reads of 1,344 transcriptomes resulted in 103,139,086 contigs, of which 163,970 were predicted to be circular. Of these putative cccRNAs, 42 were identified as viroid-like via ribozyme search (15 sequences), sequence search against ViroidDB (33 sequences), or both (6 sequences).

To verify the efficacy of the detection method, we performed a direct search of all contigs against ViroidDB and identified 12 contigs that matched a *bona fide* viroid sequence with at least 50% target coverage. The detection pipeline found four of these potentially complete viroid contigs. Of the rejected contigs, four were much larger than typical viroids (>1000 nt) and contained major ambiguous regions. The other four were low-coverage fragments that were rejected due to being smaller than unit length and therefore unable to be verified as circular. Iresine viroid 1, Citrus exocortis viroid, and a Coleus blumei viroid (CbVd) were successfully retrieved. While the first two viroids were nearly identical to the corresponding reference sequences, the CbVd-like sequence was not. At 350 nt, this sequence differed in length from all known coleviroids (CbVd 1-6 are 248-51, 301, 361-364, 295, 274, and 340-343 nt long, respectively) and in the terminal conserved region, which was identical to that of Dahlia latent viroid, suggesting an origin of this novel viroid by recombination, as has been reported for other CbVd species (Hou et al., 2009; Nie and Singh, 2017). At 85% identity, this CbVd-like sequence falls below the species membership threshold for coleviroids (Chiumenti et al., 2021).

**Table 1:**
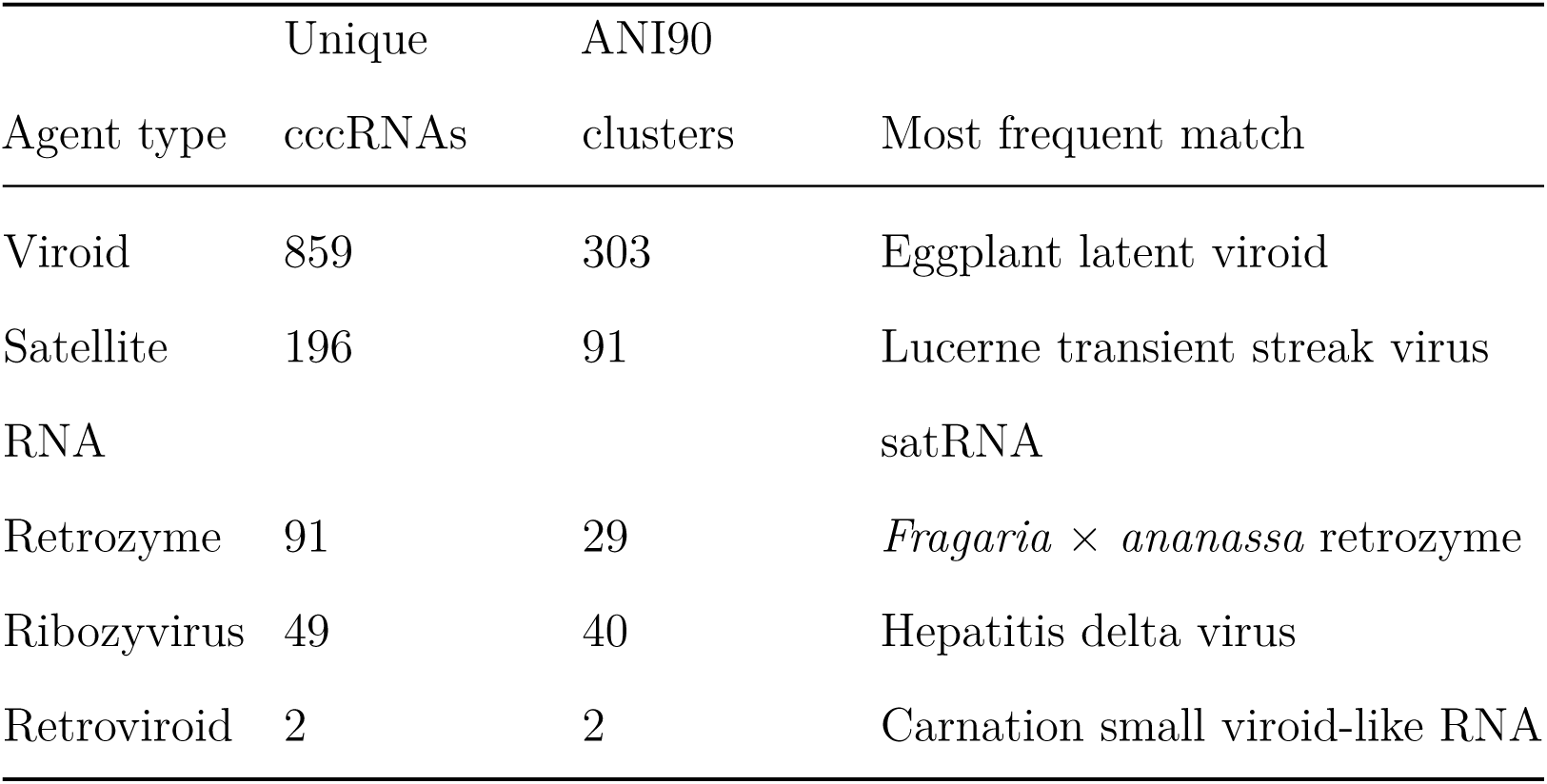
Metatranscriptomic cccRNAs and clusters with sequence matches to viroid-like agents

### A five-fold expansion of the known diversity of viroid-like cccRNAs

After testing the pipeline on plant transcriptomes, we applied it to a set of 5,131 diverse metatranscriptomes totalling 1.5 billion metatranscriptomic contigs (708 Gbp) after size filtration. The pipeline processed the entire set of transcriptomes using 720 CPU hours. We identified 10,183,455 putative cccRNAs with a median contig length of 269 nt. After removing overlapping regions and eliminating rotationally identical sequences, the median length of the 8,748,001 resultant monomers was 165 nt. Of these, 2,791,251 were within the known size range of viroids (200–400 nt), including 11,378 we classified as viroid-like because they contained a confidently predicted self-cleaving ribozyme in at least one RNA polarity. No metatranscriptomic cc-cRNAs matched the pospiviroid RY motif. Among the viroid-like cccRNAs, 10,181 were detected entirely by alignment-free methods, that is, showed no detectable direct sequence similarity (including within the ribozyme regions) to known viroids. The remaining 1,197 sequences shared significant nucleotide sequence similarity with known viroid-like RNAs spanning the entire gamut from viroids to satRNAs to retrozymes (Table 1). 907 sequences were identified as viroid-like by both the ribozyme detection and alignment-based search approaches. Among the 10,181 ribozyme-containing viroid-like cccRNAs, 3,434 were symmetric, that is, contained predicted ribozymes in both polarities. Of the 5,131 metatranscriptomes searched, 1,841 contained at least one viroid-like cccRNA.

Of the sequences aligning to viroid-like agents, the majority only contained short (<40 nt) alignable regions, generally localized to the ribozyme motifs. However, 33 sequences yielded long (>100 nt) alignments. These cccRNAs aligned to Tobacco ringspot virus satRNA (satTRSV), Lucerne transient streak virus satRNA (satLTSV), citrus dwarfing viroid (CDVd), and two retrozymes. For satTRSV and satLTSV, the range of identity between the recovered cccRNAs and the reference sequence ranged 80%–98% and 81%– 99%, respectively. The match to CDVd was 80% identical to the nearest reference sequence. In all cases, the cccRNAs were similar in length and structure to the reference sequences.

We clustered the viroid-like cccRNAs identified here in order to estimate the increase in diversity compared to previously known viroid-like RNAs (Figure 2). Aligning cccRNAs poses a challenge due to the variation in the rotation of the sequences. Two identical cccRNAs could appear to have only half the bases aligning if rotated completely out of phase. Therefore, we took special care to compensate for the circularity of the sequences during the postprocessing of the pairwise nucleotide search results (see Methods). To validate our clustering method, we tested it on ViroidDB. Previously, we identified 458 clusters at the average nucleotide identity (ANI) 90% level in ViroidDB using a method that was not circular-aware (Lee et al., 2021). We identified 50 clusters in ViroidDB using the improved method, generally corresponding to individual species.

**Figure 2:**
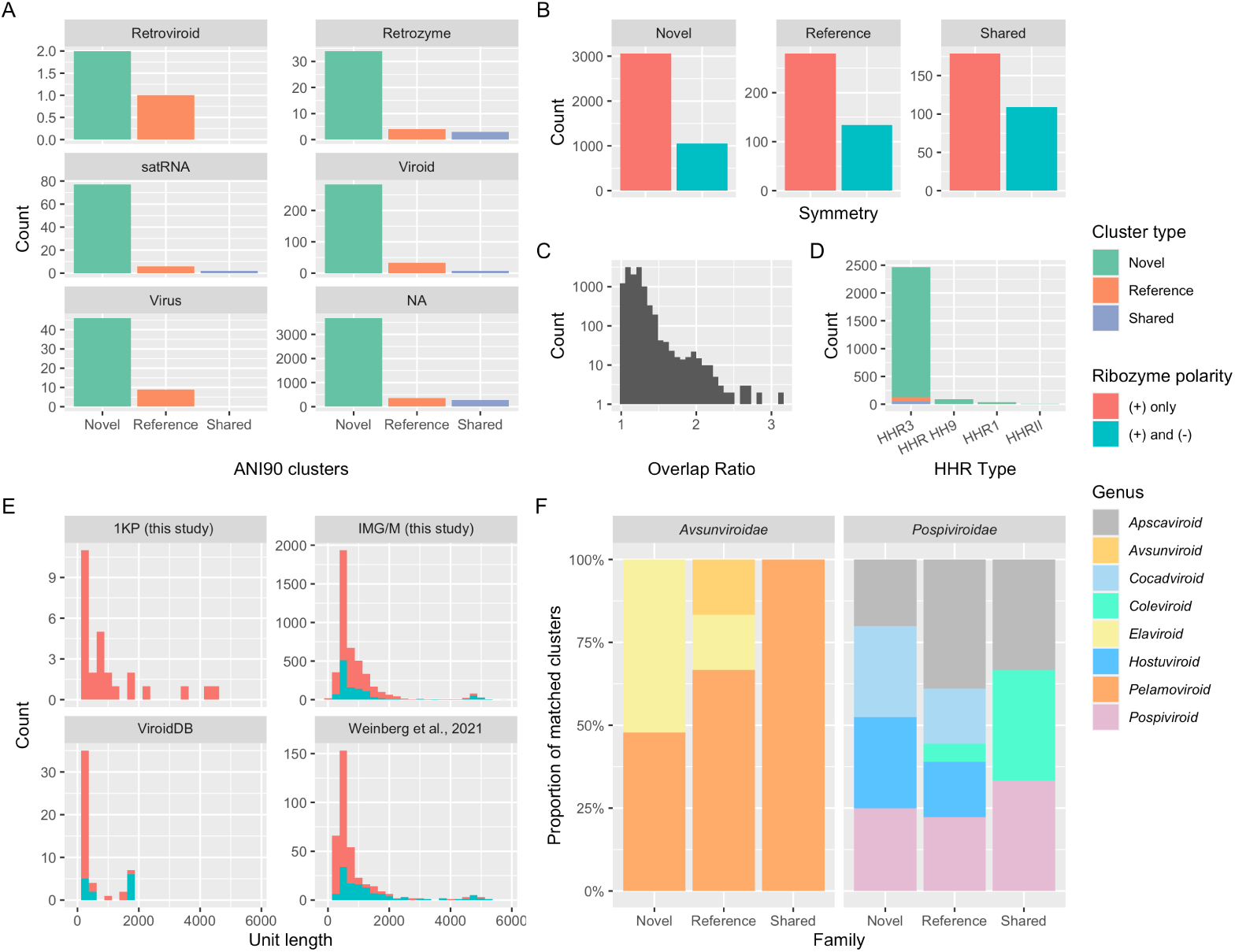
Viroid-like cccRNAs identified in metatranscriptomes. A. Number of ANI90 clusters with most significant matches to given viroid-like cccRNA agent types which are either “novel” (derived exclusively from transcriptome and metatranscriptome analysis in this work), “reference” (no novel members), or “shared” (containing at least one of both types of sequence). B. Comparative distribution of inferred ribozyme architectures by cluster type. C. Plot of overlap ratios in cccRNAs, defined as the assembled length divided by the monomer length, from IMG and 1KP. D. Counts of HHR types in representative clusters. E. Length distributions of cluster representatives in the present analysis (transcriptomes and metatranscriptomes), ViroidDB, and a previous study Weinberg et al. (2021). F. Relative abundance of clusters matching different genera within each viroid family by cluster type

In the combined metatranscriptomic, transcriptomic, and reference datasets, we identified 4,823 ANI90 clusters of which 4,121 did not include any sequences from the reference datasets and thus were considered novel, which comprises a 5.9 fold increase in viroid-like RNA diversity. Of the remaining 702 clusters containing at least one known sequence, 288 (41%) were expanded by at least one novel sequence. Notably, 39 novel clusters were represented in plant transcriptomes, of which 8 were symmetric.

The relative abundance of HHR types in the cccRNAs varied significantly from what would be expected given the sequence and species count. Within Rfam, HHR1 swamps HHR3 by two orders of magnitude by sequence count (190,679 vs 538 sequences). The same is true for the other three HHR types. However, among the cccRNA cluster representatives, the situation was inverted: HHR3 was found to be two orders of magnitude more common than HHR1 (1,952 vs 32). Similarly, HHR3 is present in less than one fifth of the HHR-containing species in Rfam whereas among the HHR-containing cluster representatives, 94% (1,952/2,074 clusters) contained at least one HHR3. Given the dominance of HHR3 in known viroids (Flores et al., 2017; Navarro et al., 2017), this overabundance of HHR3 is suggestive of the presence of numerous viroid-like cccRNAs.

### Putative novel viroid-like cccRNAs

We briefly describe the 5 largest novel ANI90 clusters (denoted 1 to 5, in the descending order of the cluster size) derived from the metatranscriptomic data to exemplify the type of findings obtained in this work. All these clusters included members with symmetric, matched ribozymes. The cccRNAs in four of these clusters contained matched HHR3s, whereas those in the fifth one contained twister-P1 ribozymes.

The largest, cluster 1, consisted of 149 sequences with a mean length of 562 nt (±9.0 nt). During circularity detection, an average of 18% of the monomer was removed, although one member of the cluster yielded a contig with 60% (341 nt) overlap. The cccRNAs in this cluster are predicted to adopt a rod-shaped conformation with 74% of the bases (on average) paired in both polarities. Most members of this cluster (*n*=137) contain symmetric HHR3s, whereas for the remaining 12 members, only one ribozyme was predicted, suggesting the presence of a divergent HHR3. Among the members of this cluster, 38 sequences yielded a short (26–37 nt) alignment to the HHR3 of Eggplant latent viroid or Grapevine hammerhead viroid-like RNA. However, the cccRNAs comprising this cluster are substantially longer than the respective viroids (334 and 375 nt, respectively). Among the members of this cluster, the majority (*n*=133) were found in terrestrial metatranscriptomes from 11 distinct locations, 4 members were identified in 3 distinct freshwater locations, and one member was found in a spruce rhizosphere sample.

The cccRNAs in cluster 2 (68 members) differed in that they contain twister-P1 ribozymes in both polarities. All but 4 of these cccRNAs are 494 nt in length and form a branched structure with a mean of 70% of self-complementary bases. These agents were found in 10 unique locations in terrestrial ecosystems, such as soil and plant litter. Nearly half (*n*=31) of the members were found in switchgrass phyllosphere samples. Cluster members were repeatedly found independently during sampling of the switchgrass phyllosphere over the course of a year at a sample site in Michigan, USA.

Like cluster 1, cluster 3 (61 members) consisted of comparatively large (605 nt), rod-shaped cccRNA with symmetric HHR3s. However, unlike the largest cluster, no member shows a detectable direct nucleotide match to any ViroidDB sequence, ribozyme or otherwise. In the majority (*n*=40) of the members symmetric ribozymes were not detected, but remaining ones were symmetric, again, suggesting the presence of divergent HHRs. Between 73% and 83% of the bases in these cccRNAs were paired in the (+) polarity and between 74% and 82% of bases were paired in the (-) polarity. On average, 28% of the monomer’s length was cleaved during circularity detection although one member was sequenced at 2.78 unit length (almost a complete head-to-tail trimer). The largest two members of the cluster (1,693 and 1,640 nt originally, 1,224 nt after cleavage) were not correctly monomerized by the circularity detection procedure due to mismatches in the seed sequence (see Methods). Manual monomerization of these sequences showed they both were 612 nt, resulting in overlap ratios of 2.76 and 2.67, respectively. Alignment results in 99.0% and 99.5% identity between monomers within each dimer, approximately the error rate of RNA polymerase II when using an RNA template (Gago et al., 2009; López-Carrasco et al., 2017; Wu and Bisaro, 2020). Almost all members of this cluster were identified in soil samples from 6 locations around the world (including Colombia, Czech Republic, Germany, and USA), and two members were found in creek freshwater samples. Cluster 4 (*n*=61) is similar to clusters 1 and 3 in that its members consist of 615 nt and are predicted to form rods containing HHR3s in both polarities.

The secondary structure results in a high mean self-complementarity of 75% of bases. However, these cccRNAs contain no alignable regions to the other large clusters or to any ViroidDB sequence. As with many other HHR3 rods, the majority (*n*=39) contain only one HHR3 above the significance threshold. All but two members of the cluster were derived from eight soil locations, and the remaining ones were found in the spruce rhizosphere.

Unique among the largest clusters we examined, the cccRNAs in cluster 5 (55 members) are quasi-rod shaped, with 64% bases paired on average in the predicted secondary structures. At 454 nt mean length (±8.9 nt), these cccRNAs are the smallest among the 5 largest clusters. In this case, all members contained two significant HHR3s, but no nucleotide matches to ViroidDB were detected. Most members were found in terrestrial samples including soil (*n*=38) and plant litter and peat (*n*=2). As in the case of cluster 2, 14 members of this cluster were also found over the span of a year in the phyllosphere of switchgrass, and one member was found in a freshwater sample. Altogether, nine distinct locations contained members of this cluster.

In summary, analysis of these 5 largest clusters showed that they consisted of cccRNAs endowed with all the hallmarks of viroids including symmetric ribozymes (HHR3s, with one notable exception), extensive (quasi) rod-shaped or branched secondary structure, and evidence of multimeric intermediates. Furthermore, the members of these clusters were independently identified in diverse samples, primarily, those from soil, indicating they are widespread and consistent with these cccRNAs being infectious agents.

### Virus-like elements blurring lines between riboviruses, ribozyviruses and viroids

Among the cccRNAs containing symmetric HHR3s, we identified rod-shaped sequences up to 4,705 nt, far outside the length range of viroids and HDVlike viruses. We hypothesized that these cccRNAs could be novel ribozy-like viruses. To perform a comprehensive search for potential ribozy-like viruses, all open reading frames (ORFs) longer than 75 codons present in cccRNAs were translated, and the resulting sequences of putative proteins were clustered by amino acid sequence similarity and compared to protein sequence databases using PSI-BLAST and HHPred (Table S2). Additionally, we searched the full set of translated cccRNAs sequences against profiles from the Pfam-A (Mistry et al., 2021) database and protein sequences from the UniRef90 (Suzek et al., 2015) database.

Notably, almost all reliable matches were to virus proteins, mostly, to capsid proteins (CP) of circoviruses, plant satellites RNA viruses, such as Satellite Tobacco Necrosis Virus, and tombus-like viruses (Table S2). One cluster of predicted proteins showed significant sequence similarity to the predicted RdRPs of a unique group of ssRNA viruses, ambiviruses, that were recently discovered in fungal isolates and transcriptomes (Forgia et al., 2021; Linnakoski et al., 2021; Sutela et al., 2020). Ambiviruses have RNA genomes of approximately 4 kb which encompass bidirectional ORFs, one of which encodes a predicted RdRP. To date, ambiviruses have not been reported to be circular (Forgia et al., 2021). In the IMG metatranscriptomes, 163 ANI90 clusters (274 cccRNAs total) were predicted to encode ambivirus-like RdRP (E-values between 1.3e-229 and 8.7e-04). Notably, these clusters of cccRNAs were also predicted to contain HHR3, HPR-meta1, and CEPB3 ribozymes, including symmetric sequences with different ribozymes in the two RNA polarities. Furthermore, all these sequences were predicted to adopt a rod-like structure in which the two ORFs encoding, respectively, the RdRP and an uncharacterized protein are arranged along the rod in the opposite strands (Figure 4). These sequences showed varying degrees of terminal overlap, with a median trimmed repeat length of 123 nt. Three representative sequences were recovered with >2000 nt overlaps, of which one was an almost-complete dimer, suggestive of replication via RCR. Three of the ambi-like clusters were detected at very low levels in 10 plant transcriptomes.

**Figure 3:**
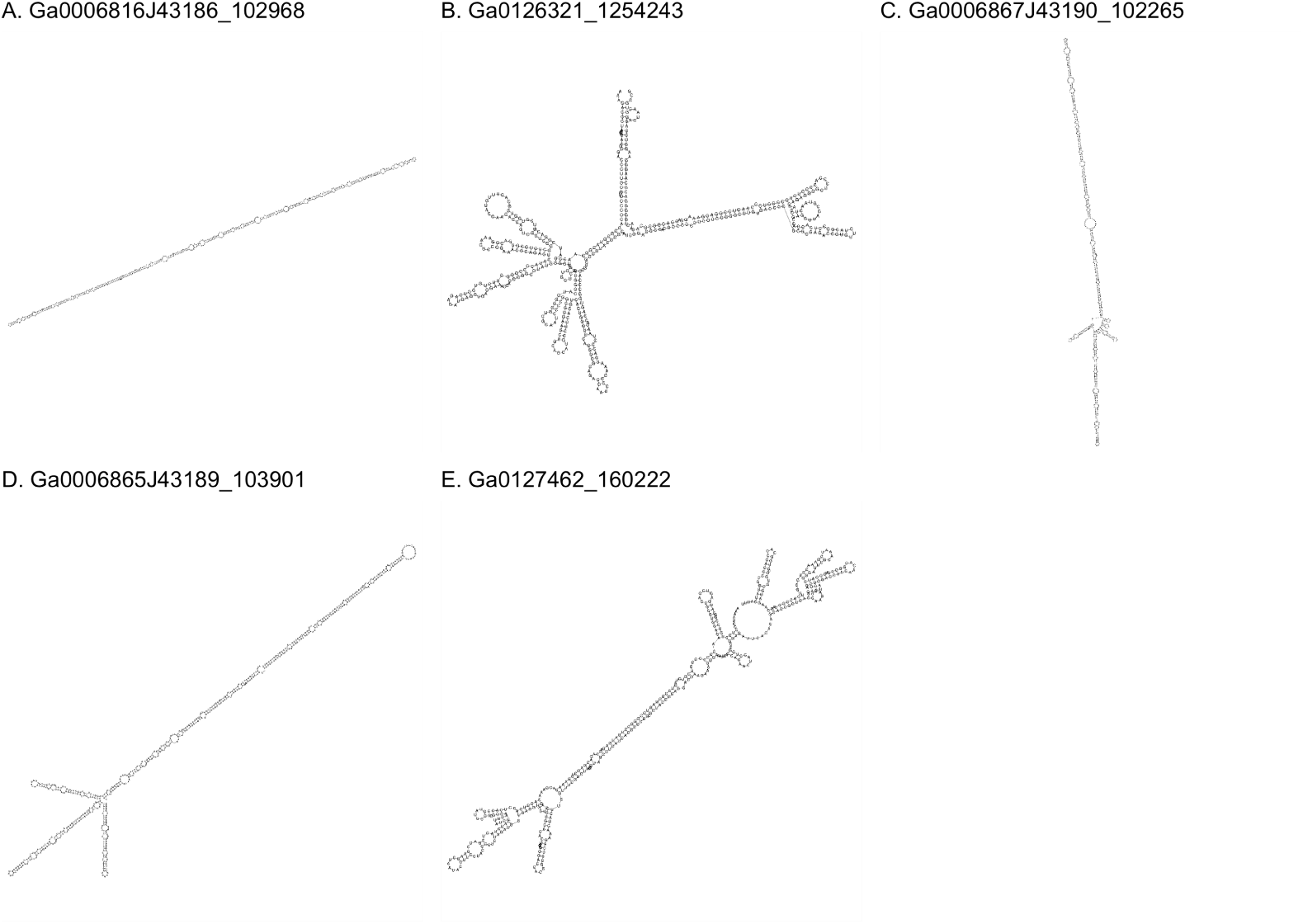
Predicted secondary structures of representatives of the five largest clusters of novel viroid-like cccRNAs. Structures were predicted using ViennaRNA’s RNAfold program configured to operate on circular sequences. Sequence data and metadata are available in Table S1.

**Figure 4:**
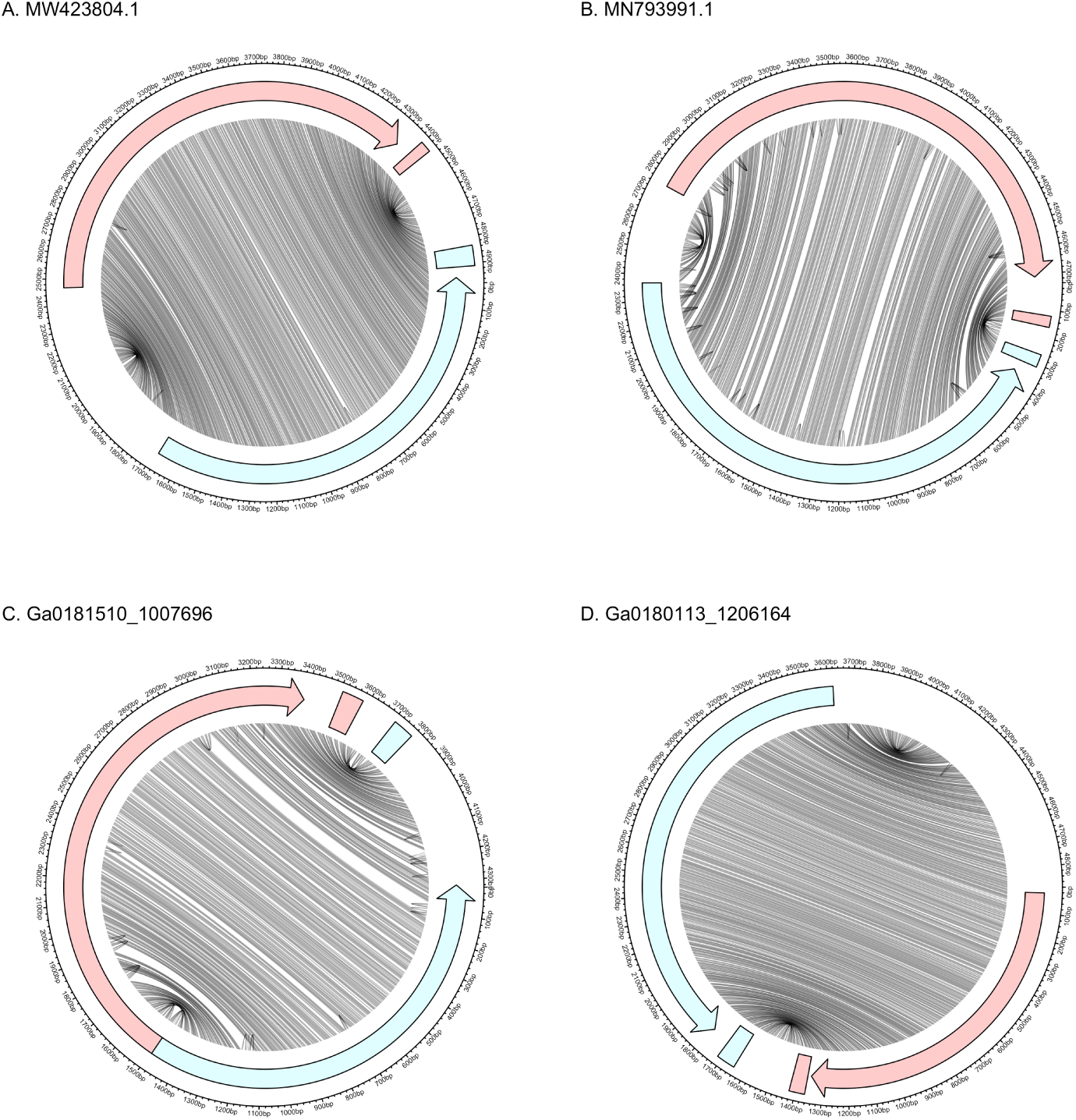
**Genomic and secondary structure** of *Armillaria borealis* ambilike virus 1 (A), *Tulasnella* ambivirus 1 (B), and novel ambi-like sequences (C, D). Red and blue denote (+) and (-) polarities, respectively. Lines connect bases in the genome that are paired in the predicted secondary structure. Arrows represent ORFs and rectangles represent self-cleaving ribozymes. The (+) and (-) ribozymes are HHR3 and HPR-meta1 in (A), HHR3 and HH3 in (B), CPEB3 and HHR3 in (C), and HHR3 and HPR-meta1 in (D). In all cases, the ribozymes are located outside the ORFs at the end of the rod.

We then ran the detection pipeline on the 30 ambivirus and ambivirus-like sequences from GenBank and detected significant ribozyme matches in 15 of these sequences, of which 13 contained two predicted ribozymes. Of the remaining 15 sequences, 11 showed ribozyme matches in the expected locations that failed to pass the significance threshold (Table S3). As in the IMG data, the HHR3 and HPR-meta1 ribozymes are present in both matched and mismatched combinations. Similarly, three of the published genomes (MT354566.2, MN793994.1, and MT354567.1) contain terminal overlaps of 160-250 nt, suggestive of circularity. Furthermore, all known ambivirus and ambivirus-like sequences were predicted to adopt a rod-shaped conformation.

Taken together, these observations strongly suggest that ambiviruses comprise a distinct group of ribozy-like viruses that encode a RdRP homologous to the RdRPs of riboviruses. There was no high similarity between the RdRP sequences of ambiviruses and any specific group of riboviruses. However, the HHPred search initiated with the alignment of ambivirus RdRP sequences yielded the top match with the RdRP profile of mitoviruses, suggesting a potential relationship with this group of capsid-less derivatives of RNA bacteriophages (leviviruses) that belongs to the phylum *Lenarviricota* in the realm *Riboviria*.

Three cccRNA clusters with significant mitovirus RdRP matches were detected, including two with symmetric ribozymes. The symmetric singletons are 3,283 and 3,058 nt in size and contain matched twister-P1 ribozymes and an HHR3/twister-P1 combination, respectively. The HHR3 aligns to ELVd with 96% identity. A third cccRNA cluster with three members encoding putative mitovirus-like RdRP, of 3,363 nt, contains a similar match to ELVd (including the HHR conserved core) that was not identified as an HHR by either detection method. This cccRNA lacks the HHR core in the opposite polarity but shows weak similarity (E-value = 0.19) to twister-P1. All cccRRNAs were detected with >100 nt overlaps. These sequences are predicted to adopt a branched conformation with between 63% and 66% of bases paired in both polarities. In all three clusters, the putative ribozymes are separated by approximately 200 nt. These three genomes have a low (36– 40%) GC content, a hallmark of mitoviruses (Marais et al., 2017). Searching the predicted RdRPs against the protein sequence databases yielded the most significant matches for the three cccRNAs to Grapevine-associated mitovirus 13, Grapevine-associated mitovirus 14, and *Fusarium asiaticum* mitovirus 8. Upon closer inspection of Grapevine-associated mitoviruses 11 and 13, we found that they also contained HHR3s in both polarities and a twister-P1/HHR3 combination, respectively. Ribozyme searches of all available *Lenarviricota* (taxid 2732407) and unclassified *Riboviria* (taxid 2585030) sequences did not yield other matches besides the ambiviruses and these few mitoviruses.

Apart from the RdRps, we identified 135 sequences comprising 53 ANI90 clusters with significant similarity to capsid proteins of single-stranded (ss) DNA viruses, in particular, CRESS viruses (Krupovic et al., 2020). The sequences in 50 of these clusters contained predicted ribozymes, and 13 contained HHR3s in both polarities. Two clusters contained paired HHR3 and twister-P1 ribozymes, whereas two other clusters contained symmetric HP-meta1 ribozymes. 26 clusters contained a single HHR3, three a twister-P1, and four a HPR-meta1. 21 clusters, including all three without ribozyme profile matches, produced a nucleotide alignment to a known viroid’s ribozyme. These alignments ranged in length from 25 to 50 nt at 83–96% identity. The cccRNAs in these clusters ranged between 1,092 and 1,632 nt in length, with a mean of 1,317 nt and GC content between 35% and 51%, with the mean of 44%. Four cccRNAs were sequenced as complete head-to-tail dimers. The secondary structure of these cccRNAs showed extensive self complementarity, with 66% of the bases paired. Given the strong evidence of circularity, extensive self-complementarity resulting in predicted branched structure and confident prediction of ribozymes, these cccRNAs most likely represent a novel class of ribozy-like satellite viruses.

### Novel ribozy-like viruses

Apart from the viruses that resembled ribozyviruses conceptually, that is, were identified as protein-coding viroid-like cccRNA but encoded proteins unrelated to HDV antigen (HDVAg), we searched for actual relatives of HDV. To this end, the metatranscriptomes were searched for cccRNAs encoding HDVAg homologs by comparing clusters of ORFs from cccRNAs to the sequences of the HDVAg and its homologs from other known ribozyviruses. A total of 12 ORF clusters were identified above the HDVAg Pfam profile’s gathering threshold; additional 21 representative ORFs were significant at the E-value < 1e-03 level, and 34 at E-value < 1e-02. Of these clusters, only one showed a significant nucleotide alignment to a known ribozyvirus. The other clusters were found in a variety of environments ranging from soil to wastewater to coastal wetland sediment. Samples with matching cccRNAs were collected from as far north as Alaska, USA, to as far south as Florida, USA. All 69 members of the clusters encompassing ORFs with HDVAg profile matches (E-value < 1e-02) were predicted to adopt a rod shape in both polarities. The genome size of ribozyviruses in ViroidDB ranges from 1,547 to 1,735 nt. However, among the HDV-like clusters identified in metatranscriptomes, the size ranged from 1,019 nt to 1,757 nt, with a median length of 1,317 nt.

Clustering of the HDVAg sequences, their homologs from other animals and the homologs from metatranscriptomes with a permissive threshold using CLANS showed that all previously known HDVAg homologs formed a single tight cluster whereas the metatranscriptome sequences formed multiple smaller clusters and singletons distant from each other and from HDV (Figure 5A). The conservation profile of the multiple alignment of the HDVAg homologs showed that the dimerization region and one of the RNA-binding regions were prominently conserved whereas the second RNA-binding region was not (Figure 5B,C). The sequences of the distant HDVAg homologs from metatranscriptomes showed low sequence similarity to the previously known HDVAgs, far below the similarity among the latter, with the distributions of percent identities almost non-overlapping (Figure 5D). Finally, in the phylogenetic tree of the HDVAg homologs, all previously known sequences formed one compact clade, whereas the homologs from metatranscriptomes identified here comprised several remaining clades, with a much greater phylogenetic depth (Figure 5E).

**Figure 5:**
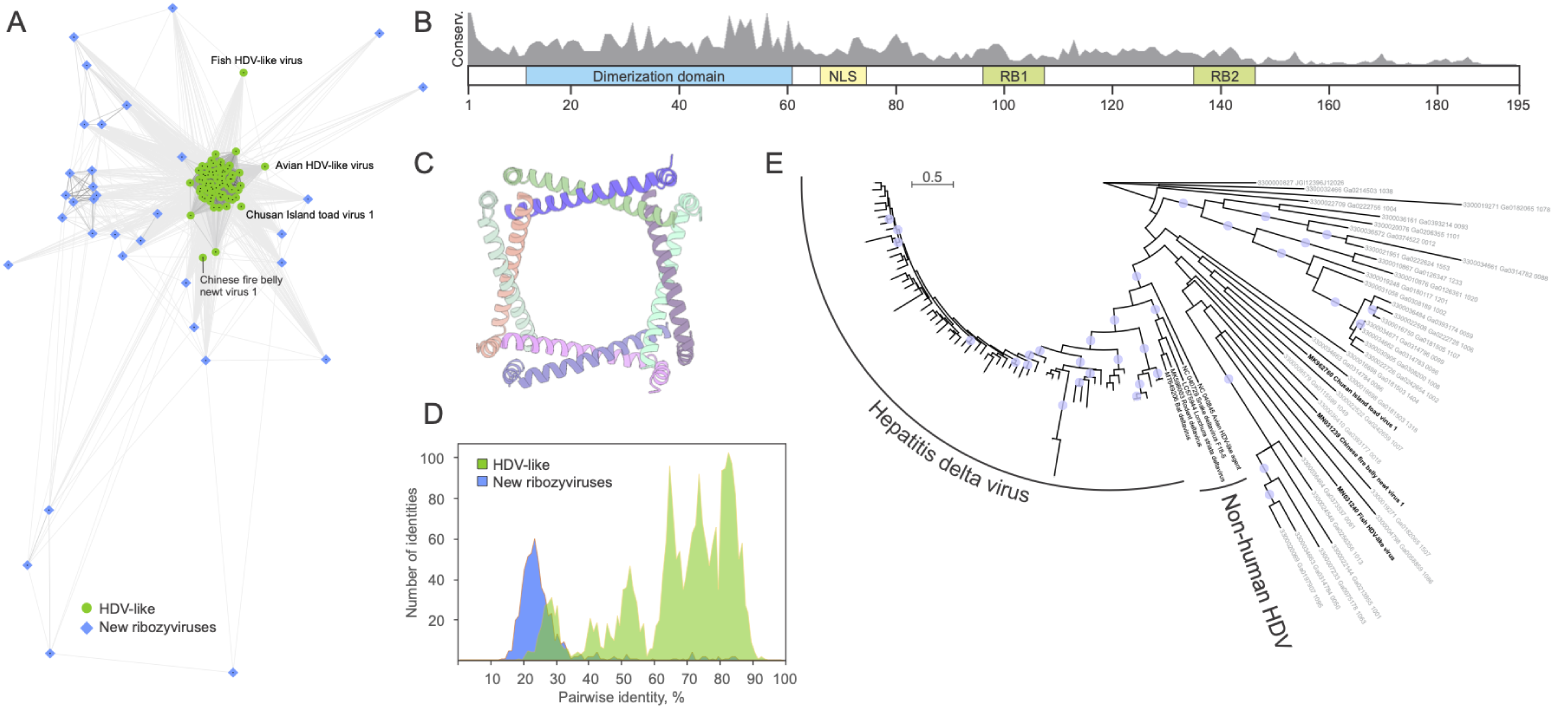
Diversity of HDV antigen-like proteins in known and newly discovered ribozyviruses. A. Clustering of the HDV antigen (Ag)-like protein homologs based on their sequence similarity. Lines connect nodes (sequences) with P-value < 1e-05. Reference HDVAg-like sequences from GenBank are shown as green circles, whereas those detected in metatranscriptomic datasets as blue diamonds. Some of the divergent reference sequences are labeled. B. Schematic representation of the HDVAg with functionally important regions indicated with colored boxes. RB1 and RB2, RNA-binding sites 1 and 2, respectively; NLS, nuclear localization signal. Gray histogram shows the sequence conservation (percent identity) of HDVAg-like sequences from metatranscriptomic datasets. C. Octameric structure of the conserved dimerization domain of HDVAg. PDB ID: 1A92 (Zuccola et al., 1998). Each protein molecule is shown with a different color. D. Comparison of the sequence conservation among reference HDVAg from GenBank (green) and those from metatranscriptomic datasets (blue). E. Maximum likelihood phylogeny of HDVAg-like sequences. The tree was constructed with IQ-TREE (Nguyen et al., 2015). Circles at the nodes represent SH-aLRT support higher than 90%. The scale bar represents the number of substitutions per site.

The nucleotide sequences of these HDV-like cccRNAs formed 26 ANI90 clusters, none which contained confidently predicted self-cleaving ribozymes above the respective gathering thresholds. However, 13 of these clusters produced weak ribozyme matches (E-value < 1e-01), and 8 of these were symmetric. Both HHR-like (*n*=36) and HDV-like (*n*=26) ribozymes were detected although no clusters contained ribozymes of both types. Of the HDV-like ribozymes detected, only five most closely matched the canonical HDV ribozyme. Ten putative ribozymes showed the strongest similarity to the HDV ribozyme (HDVR) found in the genome of *F. prausnitzii* (Webb et al., 2009), seven were most similar to the HDV-like ribozyme found in the genome *A. gambiae* (Webb et al., 2009), and four were most similar to the mammalian CPEB3 ribozyme (Chadalavada et al., 2010; Salehi-Ashtiani et al., 2006).

The limited number of significant ribozyme matches among the HDV-like sequences posed an opportunity for detecting novel ribozymes or diverged variants of known ones. For example, we examined an HDV-like cccRNA cluster (representative member 3300009579_Ga0115599_1049451) with no predicted ribozymes. However, upon closer examination, sequences from this cluster were shown to contain the conserved HHR core in both polarities in the expected locations, a recently discovered ribozyme configuration (de la Peña et al., 2021). Some clusters entirely lacked the HHR core in either polarity, suggesting the use of alternative, yet unknown ribozymes.

Clustering the HDV-like sequences in combination with the known ribozyviruses in ViroidDB produces no ANI80 clusters with both reference and novel members. Each of the 26 HDV-like clusters falls below the species demarcation criterion for ribozyviruses (80% nucleotide identity) (Hepojoki et al., 2020). Clustering the ORFs from both the detected HDV-like sequences and reference ribozyviruses with 60% minimum identity (the genus demarcation criterion) using CD-HIT resulted in 36 clusters, of which 10 consisted entirely of reference sequences whereas 26 were entirely novel.

### Novel ribozyme combinations and unusual self-cleaving ribozymes

Almost all viroid-like RNAs described to date contain the same type of ribozyme in both polarities, with the exception of some satRNAs. Surprisingly, many viroid-like cccRNAs identified in this work were predicted to contain self-cleaving ribozyme combinations that have not been so far reported in replicating cccRNAs (Figure 6).

**Figure 6:**
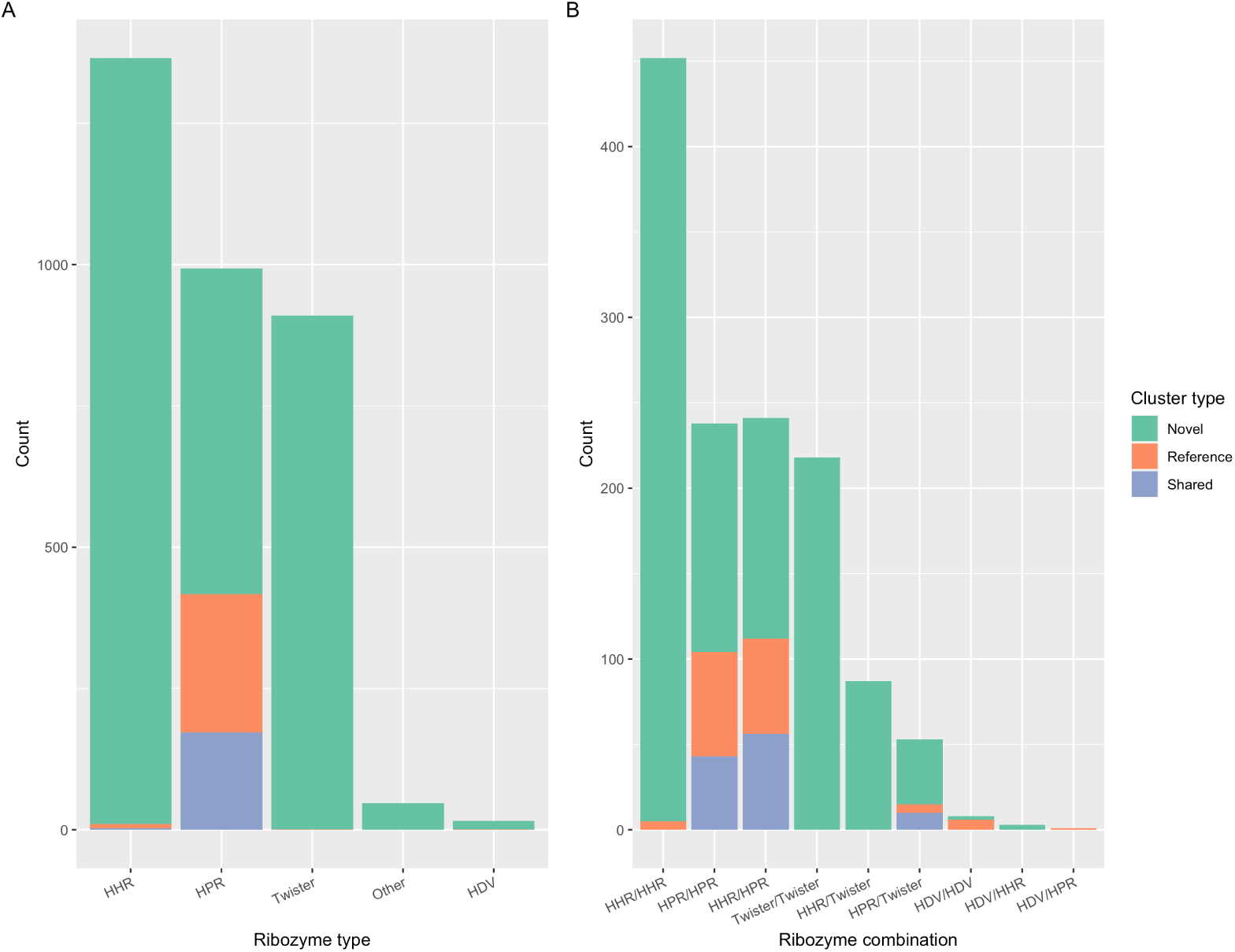
Ribozyme diversity in viroid-like cccRNAs. A. Distribution of ribozyme types in asymmetric clusters. B. Ribozyme co-occurrence within the symmetric viroid-like cccRNA cluster representatives derived from metatranscriptomes.

Specifically, we identified numerous cccRNAs containing twister ribozymes, a recently described ribozyme motif that so far has only been found in combination with the HP-meta1 ribozyme. Both symmetric (*n*=381) and asymmetric (*n*=930) variants are present in the metatranscriptomic cccRNA clusters. Most symmetric twister clusters contained matched twister ribozymes (218 clusters) in both polarities, a novel combination. In 87 clusters including mitovirus-like and satellite-like cccRNAs, we found another novel combination of ribozymes, with HHRs opposite twister ribozymes. The unusual twister ribozyme is widespread in plant transcriptomes, with 59% of the transcriptomes containing reads mapping to a twister-bearing cccRNA. Indeed, we recovered three asymmetric cccRNA clusters from plants that contained a twister-P1 ribozyme.

In addition to the twister ribozyme combinations, we identified several other novel ribozyme combinations in symmetric cccRNAs. Previously, HHR3s have been found in conjunction with HDVAg (de la Peña et al., 2021), but have not been reported to be paired with HDV ribozymes. We identified three clusters in which HHR3s were paired with HDV-type ribozymes, namely, CPEB3 and HDVR *F. prausnitzii*. The novel CPEB3-HHR3 combination was found in an ambivirus-like sequence, and the two HDVR *F. prausnitzii*/- HHR3 clusters were both predicted to adopt rod-shaped structures. One of these, a 1,052 nt singleton, did not match any sequences in ViroidDB, nt, or UniRef90, but in the other cluster (978 nt, two members), the HHR was closely similar to that of *Cryphonectria parasitica* ambivirus 1 (44/50 nt identical), whereas the HDVR *F. prausnitzii* motif (32/33 nt and 41/46 nt) aligned to two chromosomes of the *Vanessa atalanta* butterfly.

Among the asymmetric cccRNAs, we identified two additional types of self-cleaving ribozymes that so far have not been reported among viroid-like RNAs. The hatchet ribozyme was found in 34 ANI90 clusters that ranged from 357 nt to 567 nt in length and came primarily from aquatic metatran-scriptomes, in contrast to the general trend among the cccRNAs that derived from soil metatranscriptomes (Figure 7B). For example, the most diverse of these clusters (440 nt) contained 22 members from aquatic (almost all fresh-water) metatranscriptomes sampled from around the United States. For the hatchet clusters, the Rfam profile matches were the strongest among all detected ribozymes (median E-value 1.4e-08) and, unusually for viroid-like cc-cRNAs, the GC content was low (median 35%). Among ViroidDB sequences, only avocado sunblotch viroid and one retrozyme are below 40% GC. The predicted structures of these sequences varied from branched to quasi-rod shaped, with a mean of 62% of the bases paired.

**Figure 7:**
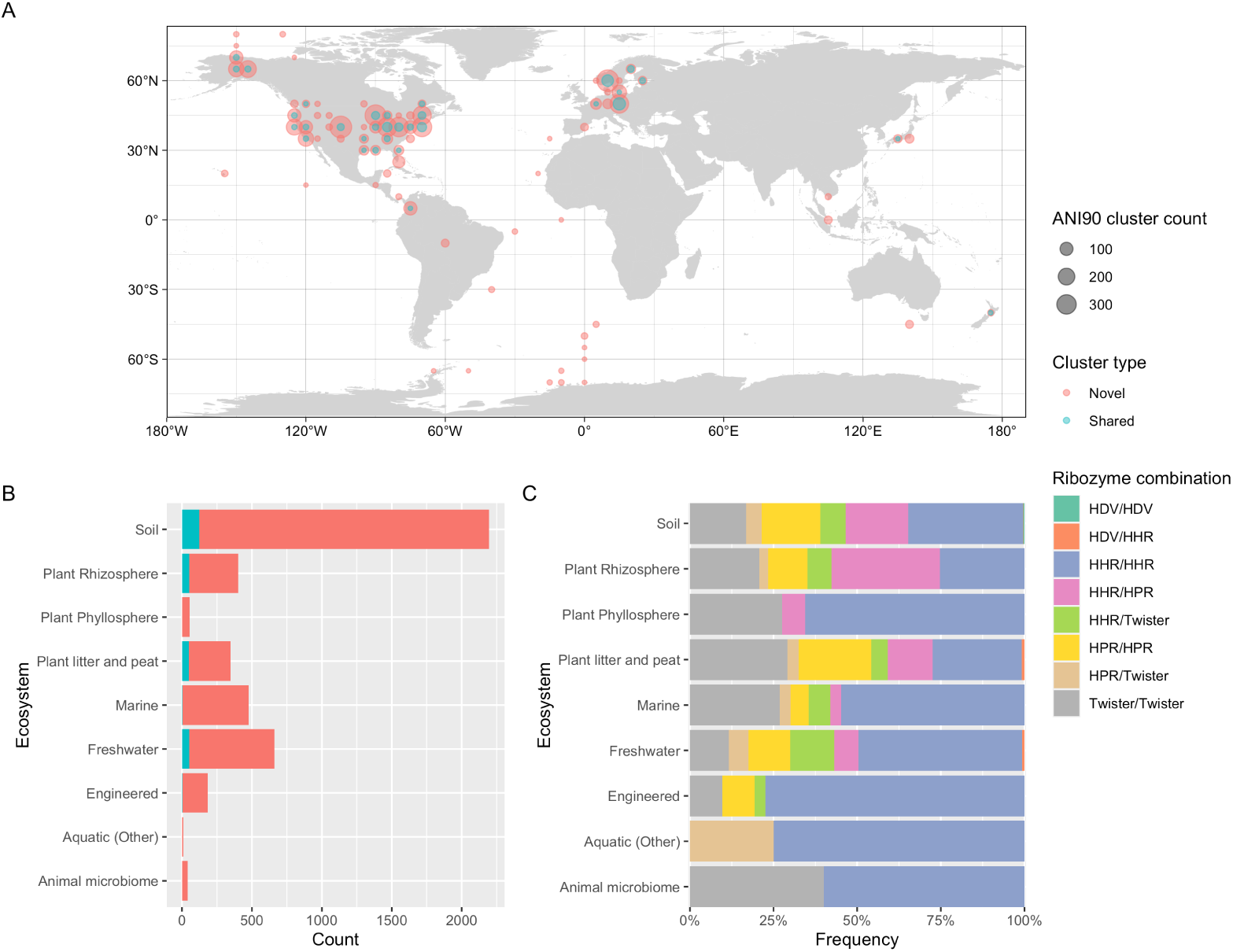
Global distribution and habitats of viroid-like cccRNAs found in metatranscriptomes. A. Map of sample locations from which viroid-like cccRNAs were detected. The size of each circle corresponds to the number of unique clusters identified in each location (grouped to the nearest five degrees of latitude and longitude) while the color represents the novelty of the clusters. B. The distribution of clusters within each type of ecosystem by novelty. C. The relative frequency of ribozyme combinations within symmetric cccRNA clusters in each ecosystem type.

The other self-cleaving ribozyme previously not known to exist in viroid-like agents, the pistol ribozyme, was identified in asymmetric cccRNAs. Like the clusters containing the hatchet ribozyme, clusters with the pistol ribozyme were found primarily (9/13 clusters) in marine metatranscriptomes ranging from the Antarctic Ocean to the Baltic Sea. The clusters have a slightly lower median profile match E-value (E-value = 6.4e-05) compared to the hatchet ribozyme but, unlike the hatchet ribozyme, have GC content ranging from 33% to 59% (median 49%) more characteristic of viroid-like RNAs. The predicted secondary structures of the pistol-containing cccRNAs were branched, often with several long hairpin structures. At least one pistol-containing cccRNA (E-value = 4.3e-11) was actually a genomic segment from a bequatrovirus (a dsDNA bacteriophage) with a short terminal repeat. The presence of a ribozyme in a bequatrovirus is surprising and represents the first report of a self-cleaving ribozyme within this genus as well as the first report of a pistol ribozyme in a virus. Further investigation is required to determine whether this virus genomic segment actually encodes a viroid-like cccRNA.

### CRISPR spacers matching cccRNAs

CRISPR spacer matches provide one of the most reliable means for assigning hosts to viruses and other mobile genetic elements in prokaryotes, and to differentiate prokaryote-infecting from eukaryote-infecting viruses (Munson-McGee et al., 2018; Paez-Espino et al., 2016). For instance, our recent search for riboviruses in the same set of metatranscriptomes that is analyzed here identified multiple spacers matching RNA viruses, resulting in the assignment of several groups of viruses to bacterial hosts including several previously thought to infect eukaryotes (Neri et al., 2022). To identify viroid-like agents that potentially might replicate in prokaryotes, we searched the viroid-like cccRNA sequences identified here against the IMG CRISPR spacer database, built from CRISPR arrays predicted in IMG genomes and metagenomes (Chen et al., 2021), and detected 89 spacers with significant matches to viroid-like cccRNAs from 9 clusters.

One spacer was an identical match of 37 nt to a member of a cluster of 16 cccRNAs with prominent viroid-like features. The cccRNAs of this cluster are 315 nt long, contain symmetric HHR3s, and are predicted to adopt a rod shape, with 73% of the bases paired. Unusual for viroids, the cccR-NAs comprising this cluster were found in Mushroom Spring, a hot spring at Yellowstone National Park. The spacer match was also detected in a 60° C hot spring, Great Boiling Spring, albeit more than 800 km away, in northwestern Nevada (Thomas et al., 2019). The repeats in this CRISPR locus were identical to those in the type III CRISPR locus of *Roseiflexus* sp. RS-1, an anoxygenic filamentous bacterium of the phylum *Chloroflexota* that was itself identified in Yellowstone hot springs (Madigan et al., 2005; van der Meer et al., 2010). Searching the same IMG metagenomic spacer database for matches to all 16 cluster members with more relaxed criteria (*i.e.*, more than 1 mismatch but with E-value < 1e-05), we identified a further 13 nearly identical matching spacers from 8 Yellowstone hot springs samples collected between 2007 and 2017. One spacer (35/37 identities, E-value = 3e-06) included a precise match to the HHR core motif. The repeats from this expanded set of loci all matched those from *Roseiflexus* sp. RS-1. In our previous study, we identified multiple spacers in *Roseiflexus* sp. RS-1 that matched a group of partiti-like riboviruses that were accordingly assigned to this bacterial host (Neri et al., 2022). Apparently, the type III CRISPR system of *Roseiflexus* sp. that encompasses a reverse transcriptase actively incorporates spacers from multiple RNA replicons.

The cluster with the most spacer matches—57 matches spanning 26 metagenomes, largely from sludge and bioreactor samples—includes a single cccRNA of 606 nt (recovered as 841 nt) with an asymmetric twister-P5 ribozyme. Eight spacers, detected in 7 metagenomes, covered the predicted ribozyme region. This sequence was predicted to adopt a branched conformation with 63% of bases paired. Nucleotide and translated nucleotide searches did not produce any matches to this cccRNA.

The second most frequently matched cluster, also a singleton, was associated with 13 spacers from 10 metagenomes, all from the same location in the Southern Indian Ocean. This cccRNA is 286 nt long (recovered with a 123 nt overlap) and contains a predicted HHRII in one polarity only. Like the most matched singleton, this sequence also is predicted to adopt a branched conformation, with 57% of bases paired. The spacers collectively covered 33% of the sequence but do not include the HHR region. No homologs of this sequence were detected in nucleotide or protein databases.

We also identified a 29 nt spacer that was identical to the HHR3 core of two distinct asymmetric cccRNA clusters. One of these is a 1,121 nt singleton and the other is a set of three 535 nt cccRNAs; both are predicted to form a branched conformation, with 68% and 65% of the bases paired, respectively. All four cccRNAs in these clusters also matched the HHR3 of Velvet mottle tobacco virus satRNA with 27/29 (93%) and 37/43 (86%) identical nucleotides, respectively. Furthermore, an identical match to the spacer was detected in the HHR of an uncultured organism found in the cerebrospinal fluid virome of a patient with unexplained encephalitis (Jimenez et al., 2011). 26 nt subsequences of the spacer also aligned with 100% identity to Chicory yellow mottle virus satRNA and Cereal yellow dwarf virus-RPA satRNA. Four of the five genes within the spacer’s scaffold (including Cas1 and reverse transcriptase) were most similar to a type III CRISPR locus of a purple sulfur bacterium, *Thiorhodococcus drewsii*. This scaffold originated from a 29° C desert spring sample from southern Nevada, whereas the cccRNA clusters were found in peatland samples from Minnesota and in spruce litter from the Bohemian Forest, Czech Republic.

### Geographic and ecological distribution of viroid-like cc-cRNAs

We examined the global distribution of the cccRNA clusters. Novel clusters were found throughout the world (Figure 7A) and in all types of ecosystems (Figure 7B). Soil samples were the primary source of both novel and shared clusters, reflecting both the greater number of such samples (twice as many as the next most common sample type) and the apparent greater sequence diversity in soils.

Surprisingly, the viroid-like cccRNAs displayed non-uniform ribozyme distribution among ecosystems (Figure 7B). Mismatched HPR/HHR ribozymes were especially prevalent among samples from plant rhizospheres, whereas matched HHRs were significantly more abundant in engineered ecosystems, such as bioreactors, than in soil environments.

Among the 10 most geographically dispersed novel clusters (that is, clus-ters with the most members found in distinct latitudes and longitudes after rounding to the nearest degree), 8 included symmetric members, of which 6 were matched HHR3s. The other two symmetric clusters contain HPR-meta1/twister-P1 ribozymes and the two asymmetric clusters contain HHR3s. These widely dispersed clusters ranged in length from 372 nt to 1,039 nt and were predicted to adopt either a rod-like shape (the 6 HHR3 clusters) or a branched conformations.

Identifying potential hosts of the viroid-like agents within these ecosystems remains a challenge. Based on the IMG annotation pipeline, the majority of the analyzed metatranscriptomes were dominated by prokaryotic sequences (Neri et al., 2022), but still contained at least 1% of contigs affiliated to eukaryotes (Table S4). Nonetheless, 187 metatranscriptomes in which viroid-like cccRNAs were detected contained < 0.1% of eukaryote-affiliated contigs, suggesting that these elements replicate in either rare and undetected eukaryotic hosts or in some of the much more abundant prokaryotic hosts. Notable among these datasets were the hot spring metatranscriptomes in which putative CRISPR targeting of viroid-like cccRNAs were identified (see above). The apparent lack of eukaryotic RNA in these samples strengthen the hypothesis of a prokaryotic host. Similarly, we found 104 symmetric clusters in marine samples which are far beyond the habitation range of the known hosts of viroids and satRNAs. To date, only ribozyviruses have been reported in aquatic hosts (Chang et al., 2019) although it has to be taken into account that viroids are highly stable and have been reported to remain infectious even after up to 7 weeks in the water (Mehle et al., 2014). The existence of these viroid-like clusters combined with the clusters from prokaryote-dominated samples suggests that the actual ecological and host range of viroid-like agents is far broader than currently appreciated.

## Discussion

Viroids and viroid-like cccRNAs, such as satRNAs and ribozyviruses, are the smallest, simplest known replicators that hijack either a host DNA-dependent RNA polymerase or a virus RdRP for their replication. Given the universality of the cellular transcription machinery across all life forms and the widespread of RdRP-encoding riboviruses, the narrow diversity and host range of the known viroid-like agents appeared puzzling. We suspected that viroid-like agents actually could be far more common than presently known and, with this motivation, searched a collection of about 5,000 metatranscriptomes for viroid-like cccRNAs.

We were able to identify millions of putative cccRNAs by identifying signatures of circularity or RCR, namely, the presence of head-to-tail repeats in assembled contigs. Because reads spanning the origin cannot be reconciled with a linear sequence, the assembler produces contigs with the same subsequence repeated at both the end and the beginning (Qin et al., 2020). Alternatively, when linear replication intermediates containing head-to-tail repeats are sequenced, the ends of the sequences are also repeated. After compensating for the low-fidelity of RNA polymerase II by allowing for up to 5% mismatches in the repeated regions, testing this method on assembled plant transcriptomes demonstrated that known viroids were reliably recovered in the absence of major assembly errors. However, the secondary structure of many viroid-like cccRNAs and the use of poly-A enrichment during RNA isolation prior to sequencing (Johnson et al., 2012) makes it likely that many viroid-like cccRNAs either were not sequenced at all or were grossly missambled and thus could not be recognized as circular. Even under a conservative approach, where only predicted cccRNAs containing confidently identified ribozymes counted as “viroid-like”, this search resulted in an approximately five-fold increase in the diversity of viroid-like agents. This is most likely a substantial underestimate of the true span of the viroid-like domain of the replicator space because among the millions of the predicted cccRNAs, in which no ribozymes were confidently identified, some, and possibly, many could be viroid-like agents containing novel ribozymes or lacking ribozymes altogether like pospiviroids. Although perhaps only a coincidence, it is worth noting that a recent analysis of the same collection of metatranscriptomes also yielded an approximately five-fold increase in the diversity of riboviruses (Neri et al., 2022).

The majority of the detected viroid-like cccRNAs possessed characteristic features of viroids including the presence of HHR, often in both polarities, and predicted rod-like or extensive branched conformation. However, the search resulted not only in quantitative but also in considerable qualitative novelty, in particular, with regard to previously undetected ribozyme combinations, such as those including twister, and detection of ribozymes not previously known to exist in viroid-like RNAs, hatchet and pistol. There seems to be little doubt that additional ribozyme combinations and novel ribozymes in viroid-like cccRNAs remain to be discovered.

Another key finding is the discovery of novel groups of ribozy-like viruses. The substantial expansion of ribozyviruses themselves, that is, viroid-like cc-cRNAs encoding homologs of the HDV antigen, probably, should have been expected. It is nevertheless notable that the diversity of the ribozyvirus sequences discovered in metatranscriptomes far exceeds that of the previously known HDV relatives including recently discovered non-mammalian ones. Moreover, many of the newly identified ribozyviruses lack the HDV ribozyme or even any known ribozymes, suggesting novel replication mechanisms. The host range of the new ribozyviruses remains to be explored but, probably, includes non-animal hosts (see discussion below).

In contrast, the demonstration that ambiviruses are actually viroid-like agents and the discovery of viroid-like mitoviruses and satellite viruses was surprising. These findings amend our understanding of the relationships between viroids and riboviruses. The latter three groups of viroid-like agents resemble ribozyviruses in that these are relatively large, protein-coding viroid-like cccRNAs. However, unlike HDV and its relatives, these viroid-like agents are clearly linked to riboviruses through the RdRPs encoded by ambiviruses and mitoviruses, and capsid proteins encoded by satellite viruses. These findings show that combinations of viroid-like cccRNA and protein-coding genes emerged on multiple occasions during evolution. It appears likely that these ribozy-like (but unrelated to HDV) viruses originated through recombination between typical riboviruses and viroids. The implications for virus taxonomy, in particular, whether such viruses should be classified into the existing realm *Ribozyviria* or into the respective divisions of the realm *Riboviria* (Koonin et al., 2020), or into completely new taxa, remain to be sorted out.

One of the most interesting but also most challenging problems is the host range of the expanded diversity of viroid-like agents. There is currently no direct computational approach for connecting viroid-like RNA with specific hosts. Nevertheless, it appears exceedingly unlikely that all or even the majority of the viroid-like cccRNAs discovered in metatranscriptomes are parasites of plants. Indeed, firstly, we identified orders of magnitude more viroid-like cccRNAs in metatranscriptomes than in plant transcriptomes, and secondly, most of the analyzed metatranscriptomes are dominated by bacteria followed by unicellular eukaryotes. Furthermore, ambiviruses were isolated from fungi (Forgia et al., 2021; Linnakoski et al., 2021; Sutela et al., 2020), and the demonstration that these are viroid-like agents expands the host range of the latter. A potential prokaryotic connection of viroid-like cccRNAs through CRISPR spacer matches is of special interest given that so far the host range of such agents appeared to be exclusively eukaryotic.

The detected spacer matches were not numerous but reliable, in particular, because multiple spacers matching the same cccRNA were identified in diverse metagenomes. At least the typical viroid-like cccRNAs that matched spacers from the reverse-transcribing type III CRISPR system of *Roseiflexus* sp. appear to be strong candidates for novel bacterial parasites. These and other viroid-like cccRNAs matched by CRISPR spacers seem to merit further, dedicated metatranscriptome and metagenome searches as well as experimental investigation. These findings, even if preliminary, echo the recent expansion of the bacterial RNA virome through the search of the same meta-transcriptome collection and suggest that bacteria might support a much greater diversity of RNA replicators than previously suspected (Neri et al., 2022).

### Limitations of the Study

This work deals with a relatively low hanging fruit in the search for viroid-like agents, being limited to the cccRNAs that contain reliably identifiable, known ribozymes or align directly to known viroid-like agents. This conservative approach was adopted purposefully, in order to avoid potential artifacts resulting from erroneous identification of cccRNA, contamination with DNA-encoded sequences or other sources. A potential opportunity for the discovery of a far greater diversity of viroid-like agents and a challenge for further research is a comprehensive analysis of the massive set of predicted cccRNAs that lack known ribozymes. Evidently, the computational approaches applied in this work only identify candidates for viroid-like agents. Experimental validation is needed and is especially important in the case of putative cccRNAs lacking known ribozymes.

## STAR Methods

### Resource availability Lead contact

Further information and requests for resources and data should be directed to and will be fulfilled by the lead contact, Benjamin Lee (benjamin.lee@chc h.ox.ac.uk).

#### Materials Availability

This study did not generate new unique reagents. All data used as inputs for the pipeline are listed in the “data acquisition” section. All output of analyses are listed in the “data and code availability” section below.

#### Data and code availability

- This paper analyzes existing, publicly available data. These accession numbers for the datasets are listed in the key resources table. Data generated during downstream analysis have been deposited at Zenodo and are publicly available as of the date of publication. DOIs are listed in the key resources table.
- All original code has been deposited at Zenodo and is publicly available as of the date of publication. DOIs are listed in the key resources table.
- Any additional information required to reanalyze the data reported in this paper is available from the lead contact upon request.

### Method Details

#### Data acquisition

The search for novel cccRNAs was performed on a collection of 5,131 assembled metatranscriptomes sourced from the IMG/MER database. In addition to the IMG metatranscriptomes, we searched the complete set of transcriptomes of the 1,000 Plants (1KP) project, totalling 1,344 paired-end sequencing runs. Before applying the search pipeline to 1KP, we filtered the raw reads for quality using fastp (Chen et al., 2018) and assembled them using rnaSPAdes (Bushmanova et al., 2019) using default parameters. We also included the 2021-09-07 release of ViroidDB and the set of HPR sequences identified by Weinberg et al. (2021).

#### Circularity detection

We identified cccRNAs using a modified and improved version of the reference-free and *de novo* Cirit algorithm (Gao et al., 2015; Qin et al., 2020) implemented in the Nim programming language. This method relies upon assembly errors for circular sequences resulting in terminal repeats. Detecting cccRNAs via this method requires searching forward for the last several bases of the sequence (the seed) and, if a match is found, comparing backwards to the start of the sequence. If the start and end of the sequence “overlap”, this repetitive region is then trimmed off. However, the existing implementation was unable to monomerize multimeric transcripts resulting from rolling circle replication among known viroid-like agents due to its single pass design and requirement of exact sequence identity within repetitive regions. Our reimplementation solves these problems by reiteratively attempting to monomerize putative cccRNAs while allowing for a configurable minimum identity within repeats. For this study, we required a minimum of 95% identity with no insertions or deletions within the overlapping region. In addition, the ratio of the length of the contig and computed monomers is reported. Sequences with monomer lengths below a threshold of 100 nt were excluded.

#### cccRNA deduplication

Standard approaches to sequence deduplication are insufficient for cccRNAs. Most modern approaches rely on hashing for memory efficiency. Such approaches are effective for linear sequences but circular sequences pose a challenge due to the arbitrariness of their start position. To enable deduplication of putative cccRNAs, we define a sequence’s canonical representation as the alphabetically earlier of the lexicographically minimal rotations of the sequence and its reverse complement. This approach, drawn from k-mer counting methods (Marçais and Kingsford, 2011; Melsted and Pritchard, 2011) and, if further optimized, would be able to be computed in linear time and with constant memory for even greater scalability (Duval, 1983).

#### Ribozyme-based filtering

To identify sequences likely to replicate via ribozyme-catalyzed rolling circle replication, we searched the unique cccRNAs for the presence of known self-cleaving ribozymes using Infernal (Nawrocki and Eddy, 2013). In each polarity, we identified ribozymes above Rfam’s curated gathering cutoff or with E-values < 0.1. Sequences with ribozymes in both polarities that met these criteria were considered viroid-like. Alternatively, we considered sequences as viroid-like with one ribozyme with an E-value < 0.01 or a score above the gathering cutoff. For each polarity, we considered only the most significant (by E-value) ribozyme. To identify more divergent ribozymes that were not detected using Infernal, we searched sequences containing one significant (E-value < 0.01) using RNAmotif (Macke, 2001).

#### Sequence search

We searched all unique cccRNAs against ViroidDB using MMseqs2 easy-search (version 13.45111) (Steinegger and Söding, 2017) with the highest available sensitivity (-s 7.5). For each sequence, we considered only the most significant match as determined by bit score. In addition to searching ViroidDB, we also searched the novel cccRNAs identified by metatranscriptome mining against the set of cccRNAs recovered from plant transcriptomes using the same method.

#### RNA secondary structure prediction

We predicted the secondary structures of all viroid-like cccRNAs for both polarities using the ViennaRNA package (Lorenz et al., 2011). For each predicted structure, we computed the percentage of bases paired and the number of hairpins present. We used a temperature of 25° C and the circular prediction mode.

#### Clustering

We performed several types of clustering including both alignment-based and alignment-free methods. To produce the alignment-based clustering, we first performed an all-versus-all search using MMseqs2 on the sequences of ViroidDB, the HPR dataset of Weinberg et al. (2021), and this study. For this method, each sequence was first concatenated to itself to compensate for potential variation in the sequence relative to otherwise-similar sequences due to their circular nature. Next, we executed MMseqs2 (v13.45111, `easy-search -s 7.5 --min-seq-id 0.40 --search-type 3 -e 0.001 -k 5 --max-seqs` 1000000) and computed the pairwise average nucleotide identity (ANI) between sequences by taking the alignment identity of the best hit for each pair. We computed the ANI for two self-concatenated sequences by first taking the length of the smaller sequence and dividing by two. We then cap the computed alignment length the the length of the now-monomzerized smaller sequence. The ANI is then defined as the percent identity within the aligned region times the alignment length divided by the smaller sequence monomer length. Similarly, we defined the alignment fraction as the smaller of the doubled query coverage, doubled target coverage, or one.

To cluster the viroids based on their pairwise ANI, we first build a graph by connecting pairs of sequences where the alignment covers at least 25% of the shorter sequence with 40% identity within the alignment. We then weighted the connections between the sequences by the ANI and employed the Leiden algorithm (as implemented in the igraph Python library, version 0.9.10) to delineate communities of similar sequences (Traag et al., 2019). The clustering granularity was optimized by iterating over the resolution parameter space until the difference between average intra-cluster ANI and the target ANI began to increase.

#### ORF prediction

To find ORFs present within the sequences, we used orfipy (Singh and Wurtele, 2021) configured to operate on circular sequences. Specifically, we searched sequences concatenated to themselves to ensure ORFs spanning the origin were detected. Only unique ORFs longer than 100 amino acids and using the standard genetic code were considered for each cccRNA.

#### Protein searches

We searched all viroid-like cccRNAs for matches to known proteins. The primary search method we used was by performing translated searches (BLASTX-style) against the UniRef90 protein database (Suzek et al., 2015) using MMseqs2 (Steinegger and Söding, 2017). For each cccRNA, we considered only the best match by E-value.

As a second approach, we also searched the ORFs from all cccRNAs, viroid-like or not, using HMMER (Eddy, 2011). We searched both the full Pfam-A profile database using hmmscan as well as a curated subset (the profiles for RdRP clan combined with the HDVAg profile) using hmmsearch.

#### HDV antigen analysis

Sequences were clustered using CLANS with BLASTP option (BLOSUM62 matrix, E-value cutoff of 1e-03) (Frickey and Lupas, 2004). Sequence similarity among reference HDVAg from GenBank and those from metatranscriptomic datasets was analyzed with the Sequence Demarcation Tool (Muhire et al., 2014). For phylogenetic analysis, HDVAg-like sequences were aligned using PROMALS3D (Pei and Grishin, 2014). Due to the short length of the sequences, the alignment was not further processed. Maximum likelihood phylogenetic analysis was performed using IQ-TREE (Nguyen et al., 2015). The best fitting model was selected by IQ-TREE and was VT+F+R4. The tree was visualized with iTOL (Letunic and Bork, 2021).

#### Read mapping

We used bowtie2 to perform read mapping from the 1KP transcriptomes to the entire viroid-like cccRNA data set in parallel. We configured bowtie to use its most sensitive setting (`--very-sensitive`) and ignore unaligned reads.

#### CRISPR spacer analysis

Viroid-like sequences were compared to predicted CRISPR spacers from prokaryotic (meta)genomes to identify potential cases of spacer acquisition from, and possible defense against, viroids by prokaryotes. The full set of 22,109 viroid and viroid-like sequences, including all reference and novel sequences, was compared to 1,961,109 CRISPR spacers predicted from whole genomes of bacteria and archaea (vJune2022) and 61,658,467 CRISPR spacers predicted from metagenomes in the IMG database (Chen et al., 2021) using BLASTN v2.9.0 with options `-dust no -word_size 7`. To minimize the number of false-positive hits due to low-complexity and/or repeat sequences, CRISPR spacers were excluded from this analysis if (i) they were encoded in a predicted CRISPR array including 2 spacers or less, (ii) less than 66% of the predicted repeats were 100% identical to each other, (iii) the spacers were at most 20 bp, or (iv) they included a low-complexity or repeat sequence as detected by dustmasker (v1.0.0) (Morgulis et al., 2006) (options `-window` 20 `-level` 10) or a direct repeat of at least 4 bp detected with etandem (Rice et al., 2000) (options `-minrepeat 4 -maxrepeat 15-threshold 2`). To initially link viroid-like sequences to CRISPR spacers, only hits with 0 or 1 mismatch over the entire spacer were considered (Table S5). To find additional spacer matches, we searched all members of the clusters with a spacer match against the IMG public metagenomic spacer data (dated 2022-06-18) set using IMG’s workspace BLAST with a minimum E-value of 1e-05. We also extracted the repeats matching loci using MinCED (Bland et al., 2007) and searched them against nt using BLASTN v2.13.0 (Altschul et al., 1990; Camacho et al., 2009).

#### Key Resources

All the code and data generated in this work are freely available through public portals listed below.

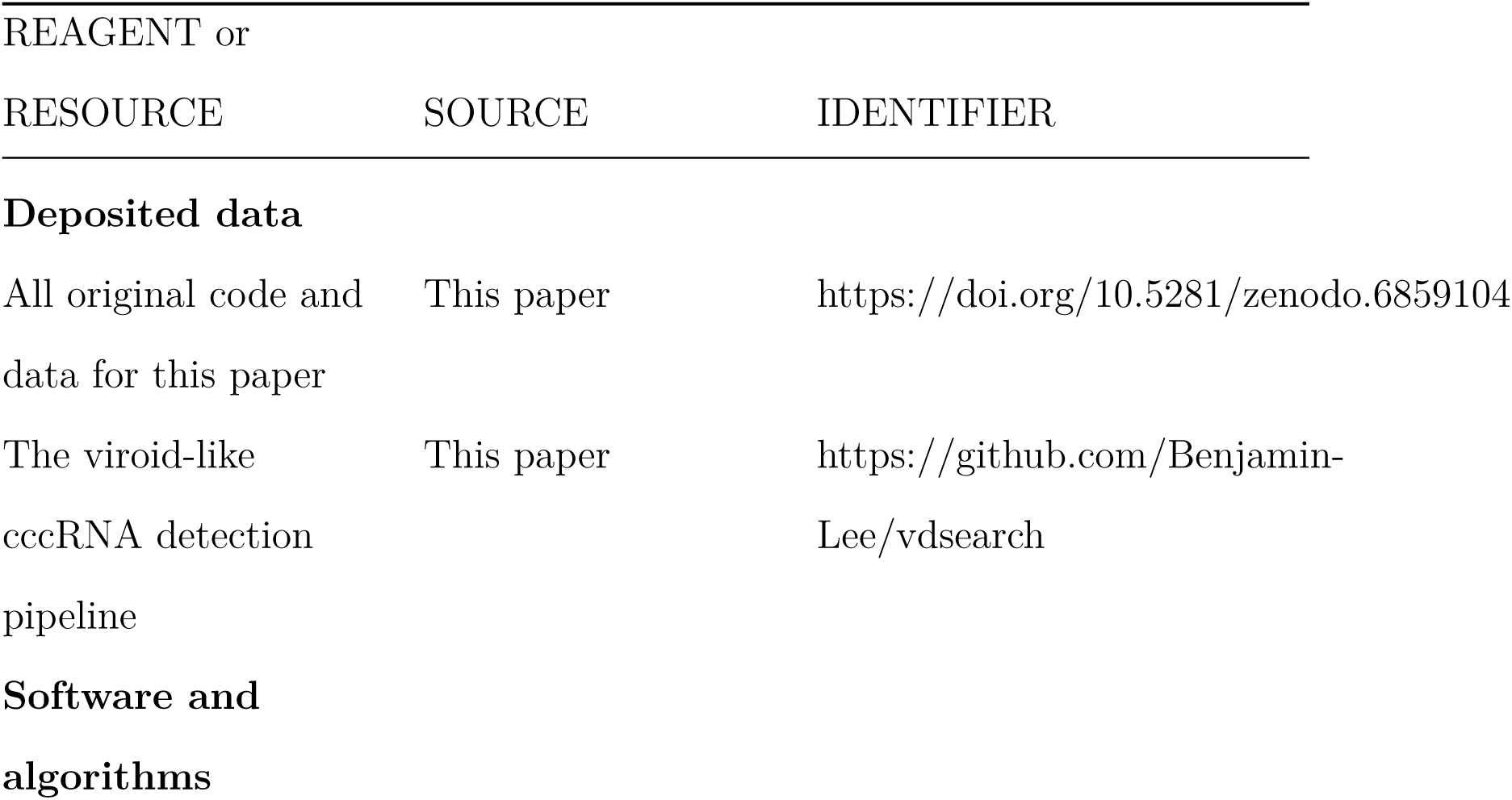
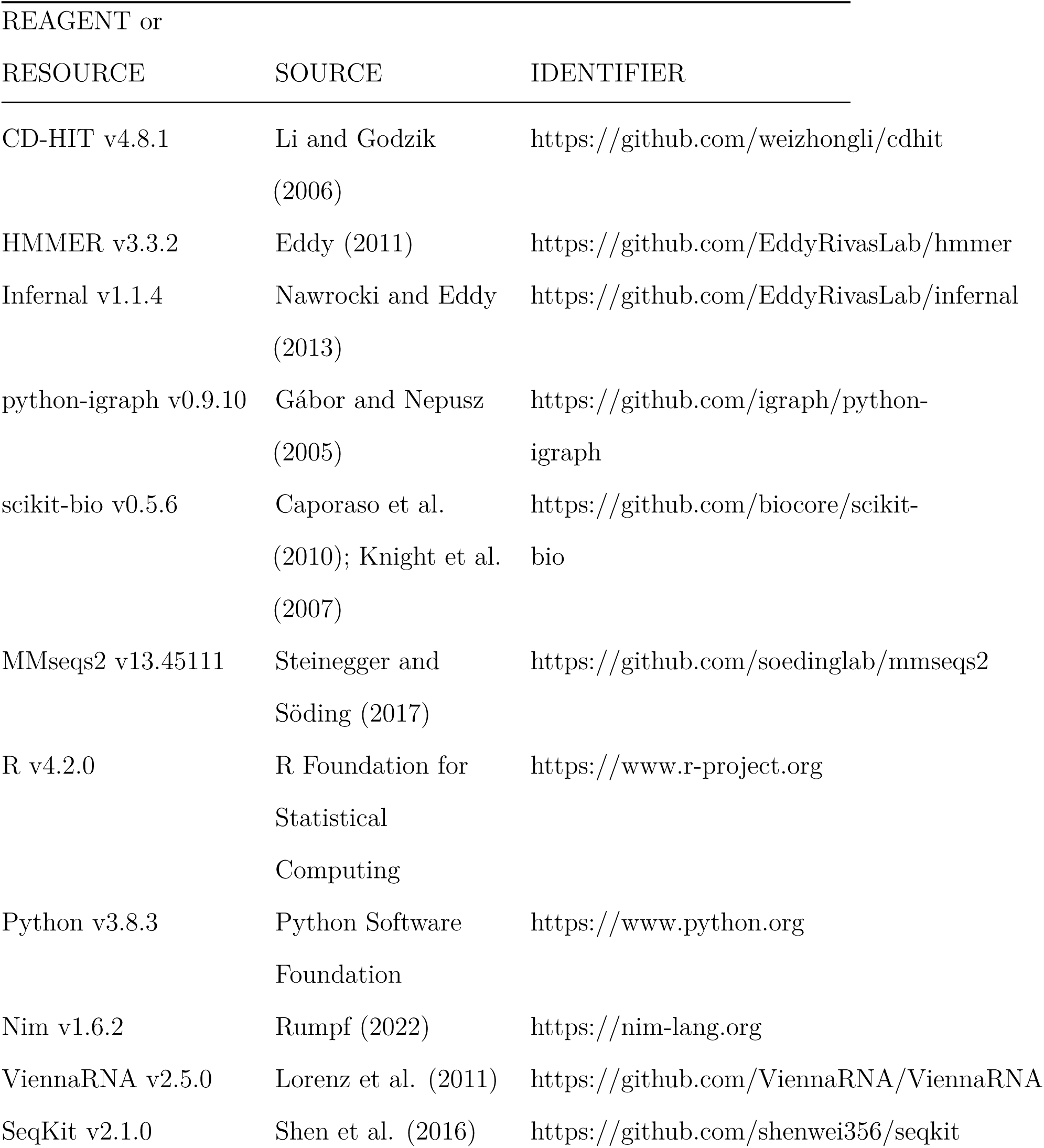
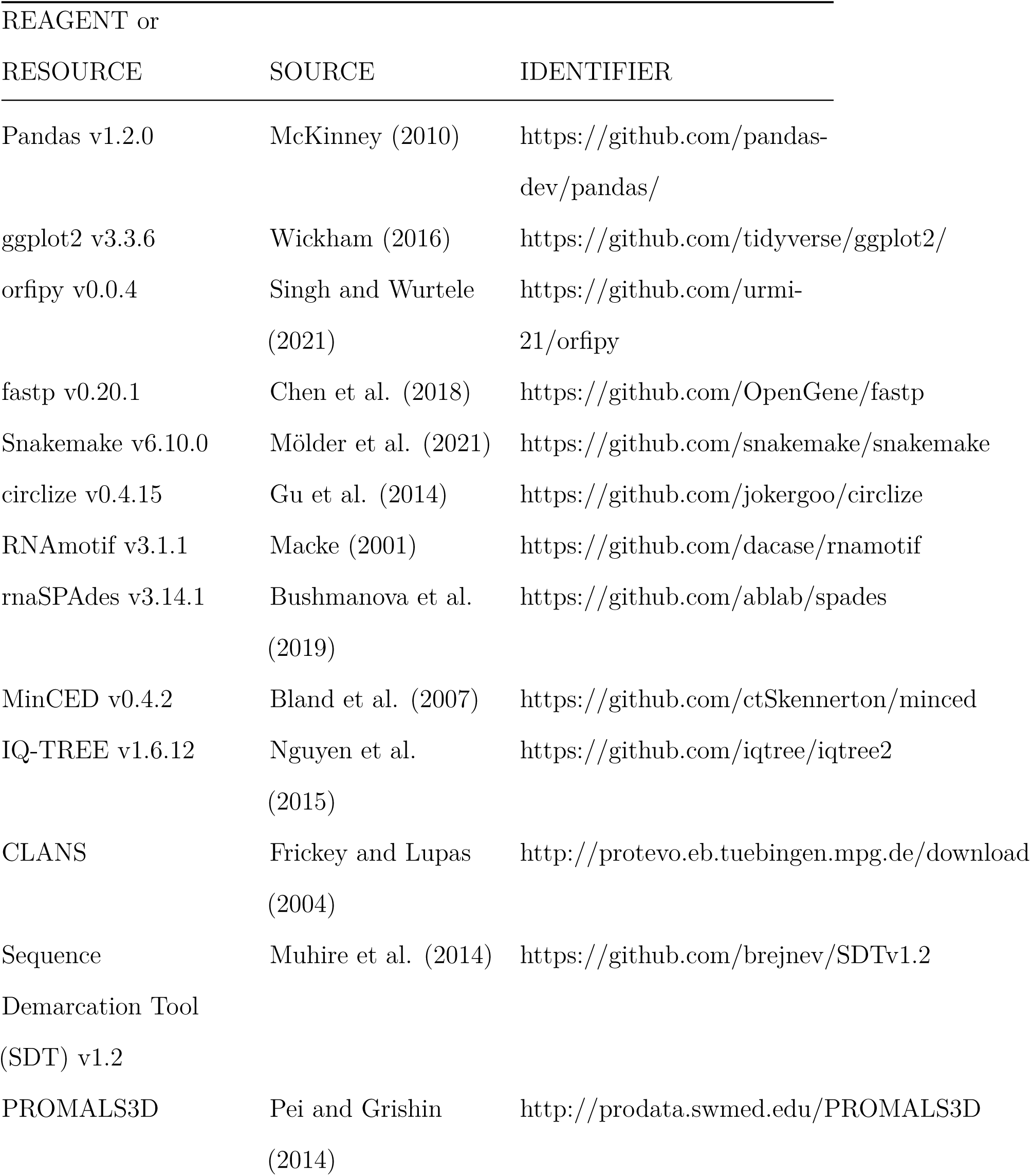
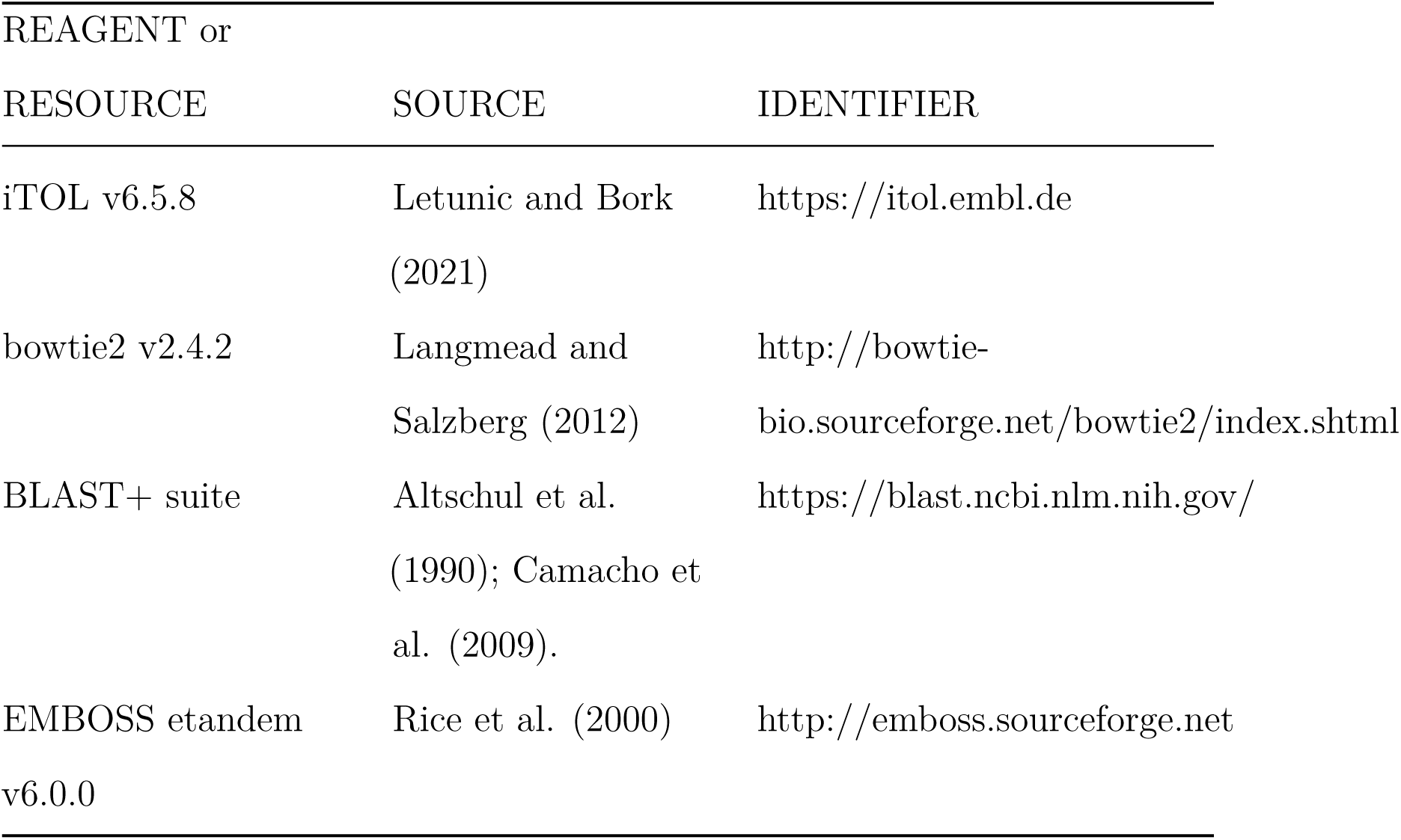

#### Deposited data

All original code and data for this paper The viroid-like cccRNA detection pipeline

### The RNA Virus Discovery Consortium

Adrienne B. Narrowe, Alexander J. Probst, Alexander Sczyrba, Annegret Kohler, Armand Séguin, Ashley Shade, Barbara J. Campbell, Björn D. Lindahl, Brandi Kiel Reese, Breanna M. Roque, Chris DeRito, Colin Averill, Daniel Cullen, David A. C. Beck, David A. Walsh, David M. Ward, Donald A. Bryant, Dongying Wu, Emiley Eloe-Fadrosh, Eoin L. Brodie, Erica B. Young, Erik A. Lilleskov, Federico J. Castillo, Francis M. Martin, Gary R. LeCleir, Graeme T. Attwood, Hinsby Cadillo-Quiroz, Holly M. Simon, Ian Hewson, Igor V. Grigoriev, James M. Tiedje, Janet K. Jansson, Janey Lee, Jean S. VanderGheynst, Jeff Dangl, Jeff S. Bowman, Jeffrey L. Blanchard, Jennifer L. Bowen, Jiangbing Xu, Jillian F. Banfield, Jody W Deming, Joel E. Kostka, John M. Gladden, Josephine Z Rapp, Joshua Sharpe, Katherine D. McMahon, Kathleen K. Treseder, Kay D. Bidle, Kelly C. Wrighton, Kimberlee Thamatrakoln, Klaus Nusslein, Laura K. Meredith, Lucia Ramirez, Marc Buee, Marcel Huntemann, Marina G. Kalyuzhnaya, Mark P Waldrop, Matthew B Sullivan, Matthew O. Schrenk, Matthias Hess, Michael A. Vega, Michelle A. O’Malley, Monica Medina, Naomi E. Gilbert, Nathalie Delherbe, Olivia U. Mason, Paul Dijkstra, Peter F. Chuckran, Petr Baldrian, Philippe Constant, Ramunas Stepanauskas, Rebecca A. Daly, Regina Lamendella, Robert J Gruninger, Robert M. McKay, Samuel Hylander, Sarah L. Lebeis, Sarah P Esser, Silvia G. Acinas, Steven S. Wilhelm, Steven W. Singer, Susannah S. Tringe, Tanja Woyke, TBK Reddy, Terrence H. Bell, Thomas Mock, Tim McAllister, Vera Thiel, Vincent J. Denef, Wen-Tso Liu, Willm Martens-Habbena, Xiao-Jun Allen Liu, Zachary S. Cooper, Zhong Wang

## Acknowledgements

The authors would like to thank Samuel Wilder for his support while developing the software pipeline and Caleb Oh for advice on software architecture. This work utilized the computational resources of the NIH HPC Biowulf cluster (http://hpc.nih.gov). Figure 1 was created with BioRender.com. B.D.L. was supported by a fellowship from the National Institutes of Health Oxford-Cambridge Scholars Program. Y.I.W. and E.V.K. are supported through the Intramural Research Program of the US National Institutes of Health (National Library of Medicine). U.G. and U.N. are supported by the European Research Council (ERC-AdG 787514). U.N. is partially supported by a fellowship from the Edmond J. Safra Center for Bioinformatics at Tel Aviv University. V.V.D. was partially supported by NIH/NLM/NCBI Visiting Scientist Fellowship. The work of the U.S. Department of Energy Joint Genome Institute (S.R., N.K. and all JGI co-authors), a DOE Office of Science User Facility, is supported by the Office of Science of the U.S. Department of Energy under contract no. DE-AC02-05CH11231. M.K. was supported by l’Agence Nationale de la Recherche grants ANR-20-CE20-009-02 and ANR-21-CE11-0001-01.

## Author Contributions

E.V.K. incepted the project; B.D.L. and E.V.K. designed research; B.D.L. and U.N. compiled the datasets; B.D.L, U.N., S.R., A.P.C., Y.I.W and M.K. analyzed the data; P.S., U.G., N.K., V.V.D. and E.V.K. supervised research; BDL and E.V.K. wrote the manuscript that was read, edited and approved by all authors.

## Supplementary Information

- Table S1: Summary of each viroid-like cccRNA from plant transcriptomes, metatranscriptomes, ViroidDB, and Weinberg et al. (2021)
- Table S2: Predicted proteins in cccRNAs
- Table S3: Predicted self-cleaving ribozymes in known ambi- and ambi-like viruses
- Table S4: Sample preparation method and taxonomic composition for each metatranscriptome
- Table S5: CRISPR spacer matches to cccRNAs

